# Recording morphogen signals reveals origins of gastruloid symmetry breaking

**DOI:** 10.1101/2023.06.02.543474

**Authors:** Harold M. McNamara, Sabrina C. Solley, Britt Adamson, Michelle M. Chan, Jared E. Toettcher

## Abstract

When cultured in three dimensional spheroids, mammalian stem cells can reproducibly self-organize a single anterior-posterior axis and sequentially differentiate into structures resembling the primitive streak and tailbud. Whereas the embryo’s body axes are instructed by spatially patterned extra-embryonic cues, it is unknown how these stem cell gastruloids break symmetry to reproducibly define a single anterior-posterior (A-P) axis. Here, we use synthetic gene circuits to trace how early intracellular signals predict cells’ future anterior-posterior position in the gastruloid. We show that Wnt signaling evolves from a homogeneous state to a polarized state, and identify a critical 6-hour time period when single-cell Wnt activity predicts future cellular position, prior to the appearance of polarized signaling patterns or morphology. Single-cell RNA sequencing and live-imaging reveal that early Wnt-high and Wnt-low cells contribute to distinct cell types and suggest that axial symmetry breaking is driven by sorting rearrangements involving differential cell adhesion. We further extend our approach to other canonical embryonic signaling pathways, revealing that even earlier heterogeneity in TGFβ signaling predicts A-P position and modulates Wnt signaling during the critical time period. Our study reveals a sequence of dynamic cellular processes that transform a uniform cell aggregate into a polarized structure and demonstrates that a morphological axis can emerge out of signaling heterogeneity and cell movements even in the absence of exogenous patterning cues.

**Highlights:**

- Symmetry-breaking gastruloid protocol where Wnt signaling evolves from a uniform high state to a single posterior domain.
- Synthetic gene circuits record Wnt, Nodal and BMP signaling with high time resolution.
- Heterogeneity in Wnt signaling at 96 h predicts cells’ future positions and types.
- Wnt activity differences are influenced by earlier heterogeneity in Nodal/BMP activity.

## Introduction

One of the most remarkable features of a developing embryo is its capacity for self-organization. Developmental self-organization arises in many contexts, but generally features cells generating or amplifying asymmetries to progress from a nearly uniform initial state to a strongly polarized outcome. Its importance has been appreciated for at least a century, when Spemann and Mangold demonstrated that cells transplanted from the dorsal blastopore lip of one amphibian embryo into another were sufficient to recruit host cells into forming a secondary body axis^1^. Since these foundational experiments, self-organizing potential has been demonstrated in a range of model organisms and developmental processes, and is not exclusively localized to discrete ‘organizer’ regions but can be broadly distributed throughout the embryo^2^. Understanding how embryonic cells achieve reproducible outcomes through self-organizing processes is a fundamental challenge in developmental and systems biology, and may suggest engineering routes towards regenerative medicine and biomanufacturing.

Developmental self-organization can in principle be orchestrated through many classes of molecular and physical processes. At the molecular level, diffusible ligands can spread between cells which then can respond by secreting either the same or a different ligand. Cells thereby form feedback loops that amplify small differences and generate stable patterns^3–5^. For example, injection of mRNA encoding Nodal- and BMP-family ligands into the zebrafish embryo is sufficient to induce host cells to form a complete secondary body axis^6^. A second class of mechanisms involves physical interactions between cells. For example, cells expressing different adhesion receptors can self-sort based on the relative strengths of their interactions to form stable patterns in both natural and synthetic contexts^7, 8^. These mechanisms need not be mutually exclusive; physical interactions and intracellular signaling can influence one another in mechanochemical feedback loops^9^. Due to this complexity, we lack a complete understanding of developmental self-organization in virtually every context in which it has been observed. What are the earliest signaling asymmetries that contribute to a polarized outcome, and how are they subsequently amplified and stabilized by biochemical and physical interactions between cells?

The recent emergence of organoid-based models of embryogenesis provides an opportunity to study self-organization in a simplified and well-controlled context. For example, 3-dimensional gastruloids derived from mouse or human embryonic stem cells are able to break symmetry and elongate along a single anterior-posterior body axis in response to uniform signaling stimuli (*e.g.*, a 24 h pulse of the Wnt pathway activator CHIR-99021) over an experimentally tractable timescale of approximately 5 days^10–12^. Gastruloid polarization and axial elongation resembles the formation of the embryonic primitive streak during gastrulation, where the epiblast’s radial symmetry is broken to establish the anterior-posterior body axis and form the three germ layers^13^. Whereas gastrulation *in vivo* is guided by spatially patterned external cues (e.g., Wnt3 secreted from the visceral endoderm; BMP4 from the extra-embryonic ectoderm), gastruloids organize a single body axis in the absence of any extra-embryonic tissues or asymmetrically applied cues. This capability to polarize without an apparent pre-pattern is consistent with a capacity for self-organization that is already present in the embryo, where transplantation of the anterior primitive streak (or ‘node’) is sufficient to induce duplication of a secondary neural axis^14^.

How gastruloids spontaneously break symmetry to form a polarized body axis remains an open question. Treatment with signaling inhibitors can identify signaling pathways involved in polarization^15–17^, but often produces broad defects (e.g. failure to polarize) which cannot dissect dynamics of self-organization. Current signaling biosensors are also insufficient: while prior studies have characterized spatiotemporal patterns of BMP, Nodal, and Wnt signaling using transcriptional reporters^10^, it has been challenging to causally link cells’ signaling histories to their subsequent position along the anterior-posterior (A-P) axis. Gastruloids are optically dense, can contain 10^4^-10^6^ cells, and are highly dynamic, with considerable cell migration, rearrangement, and differential proliferation^17, 18^, making imaging-based single-cell tracking difficult if not impossible using current methods. While powerful, standard transcriptomic techniques (e.g., scRNAseq; smFISH) are terminal measurements and thus also poorly suited to relate cells’ current states to their prior history, future fates, or eventual spatial position.

Here, we describe a synthetic biology approach to map how early signaling dynamics predict cells’ future spatial position in the developing gastruloid. We engineered mouse embryonic stem cells expressing “signal-recorder” gene circuits to permanently label cells in which a particular pathway was active in a user-defined time window. We show that these recombinase-based circuits can record morphogen signals with high fidelity and temporal resolution, achieving near-complete labeling of the currently signaling-active population within 6 h. By systematically varying the recording time and measuring labeled cells’ final positions, we map how signaling dynamics encode fate information over time. We first use this approach to identify the earliest time points at which Wnt activity is predictive of cells’ future A-P axis fates. We define a “symmetry-breaking” gastruloid protocol where Wnt signaling progresses sequentially through uniformly low, uniformly high, and heterogeneous states before localizing exclusively to the posterior pole. Surprisingly, we find that Wnt activity at 96 h post-seeding is already predictive of cells’ future A-P position and fate, even though neither Wnt activity nor gastruloid morphology is polarized by this time. Single-cell sequencing and live imaging measurements suggest that these early Wnt-active and Wnt-inactive populations undergo cell sorting to define anterior and posterior cell populations, likely driven by differential cell-cell adhesion. Finally, we extend our signal recording approach to Nodal and BMP signaling, revealing that even earlier pre-existing heterogeneity in both pathways is predictive of cells’ future axial positions, likely by altering subsequent Wnt activity dynamics. Taken together, our data reveal early signaling events associated with symmetry-breaking in a model of mammalian development, suggesting a model where early stochastic signaling events and subsequent cell sorting are sufficient to specify the anterior-posterior axis even in the absence of spatially restricted extra-embryonic cues.

## Results

### Wnt signaling evolves dynamically during gastruloid polarization

We began our investigation of symmetry breaking by focusing on the Wnt signaling pathway, which coordinates A-P patterning in many bilaterally symmetric animals^19^. Gastruloid morphogenesis is triggered by transient activation of Wnt pathway activity via addition of the small molecule CHIR-99021 (“CHIR”) to the culture medium. A short time later, Wnt activity becomes polarized to the gastruloid’s posterior domain, where it is required for further development at the morphological and molecular level (e.g., Brachyury expression)^11, 20^. Polarized Wnt activity is thought to act in concert with FGF signaling activity to define an organizing domain capable of autonomously driving gastruloid elongation^21^. This localized posterior domain of Wnt activity mirrors the situation in the embryo, where Wnt3 is provided as a spatially restricted extra-embryonic cue by the visceral endoderm to initiate gastrulation within the posterior region of the epiblast^22^. We hypothesized that the gastruloid’s progression from a global Wnt-active state (during CHIR stimulation) to a localized posterior domain could reflect an underlying program of developmental self-organization (**Figure 1A**), so we began our study by characterizing how a polarized Wnt signaling domain emerges following a uniform stimulus.

**Figure 1:**
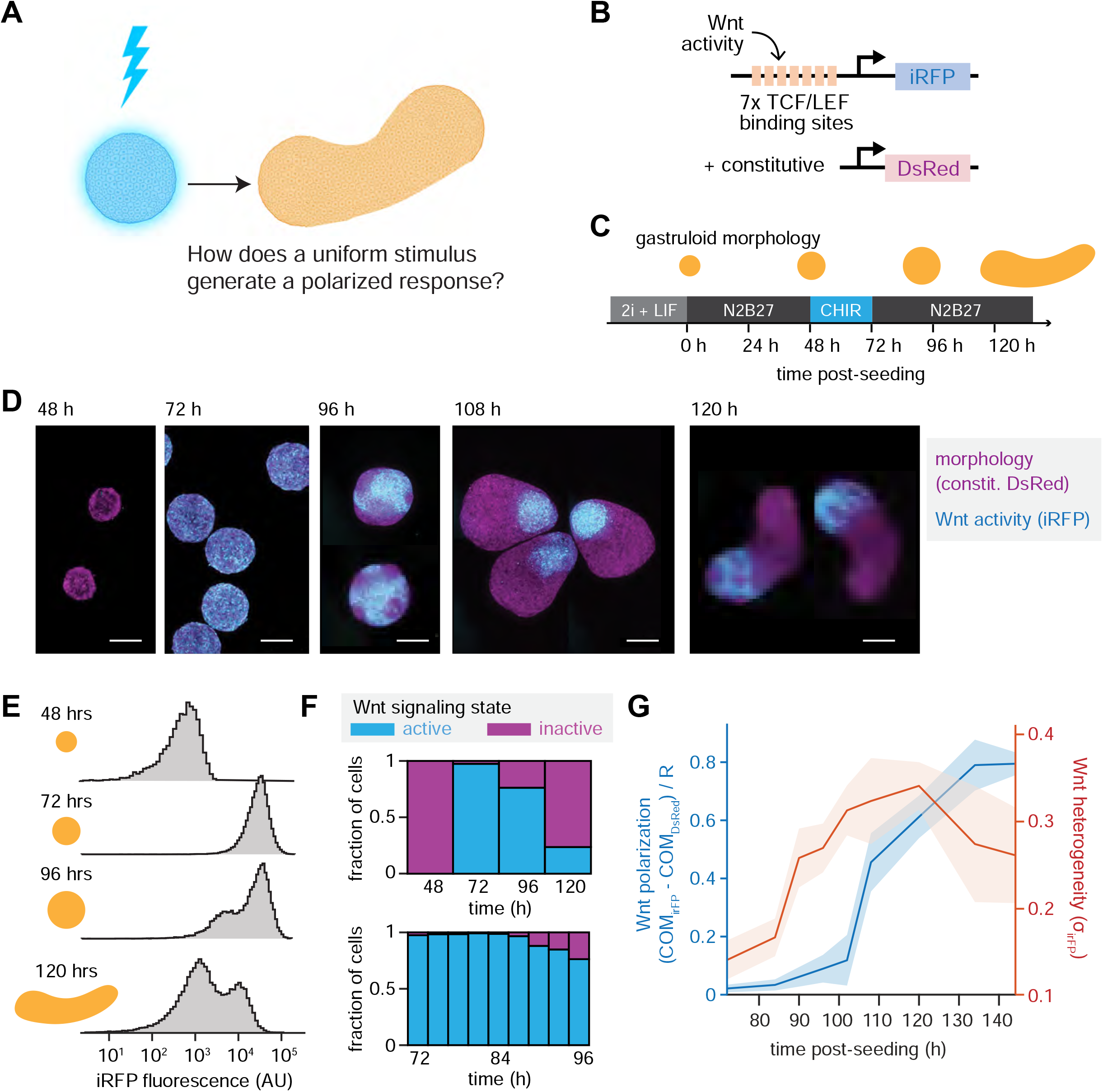
Dynamics of Wnt symmetry breaking and polarization during gastruloid morphogenesis. **(A)** Illustration of gastruloid self-organization phenomenon: a transient, spatially uniform stimulus somehow triggers the formation of a polarized morphology without an exogenous pre-pattern. **(B)** A clonal mESC line was engineered to report Wnt activity through the expression of a destabilized iRFP downstream of Wnt-sensitive TCF/LEF enhancer sites. **(C)** ‘Symmetry-breaking’ protocol for gastruloid generation. mESC cultures were maintained in 2i+LIF culture media until immediately before gastruloid formation to suppress pre-existing heterogeneity in Wnt activity. Gastruloids were formed from 200 initial cells/gastruloid seeded using a cell sorter, and treated with 3 µM CHIR between t=48 and 72 h to stimulate morphogenesis. **(D)** Dynamics of Wnt activity patterns during gastruloid morphogenesis. Samples were fixed at variable timepoints and imaged to measure spatial distributions of Wnt signaling. **(E)** Single-cell Wnt activity levels were measured by flow cytometry. Histograms of Wnt activity indicate an initially uniform response to Wnt activation with CHIR at 72 h, followed by a bimodal response. **(F)** Quantification of the proportion of cells which are Wnt active over time. A Wnt-inactive population is first detectable at t = 90 hours. **(G)** Quantification of heterogeneity and polarization in spatial patterns of Wnt activity during gastruloid morphogenesis (n = 76 gastruloids). Heterogeneity is reported as a normalized standard deviation, and polarization is reported as a normalized distance between the center of mass (COM) of Wnt activity and morphological images. Quantification indicates an onset of heterogeneity at t = 90 h (consistent with flow cytometry), followed by polarization at t = 108 h.

We engineered a clonal mESC line in which a Wnt-dependent promoter drives expression of a destabilized infrared fluorescent protein (P_TCF/LEF_-iRFP-PEST)^23^ and where constitutive DsRed expression marks all cells (**Figure 1B**). Benchmarking the dynamics of this biosensor revealed that the Wnt reporter responds within 5 h to acute changes in pathway activity (**Figure S1A**). We also optimized gastruloid culture conditions to monitor the evolution of Wnt signaling patterns from the uniform to polarized state. Gastruloids can be seeded from mESCs that were cultured in LIF-supplemented growth media or in 2i+LIF media, where the latter media contains CHIR and a MEK inhibitor to suppress spontaneous differentiation and preserve culture uniformity^24^. We found that the standard protocol, in which gastruloids are seeded from cells cultured in LIF-supplemented growth media (**Figure S1B**), led to substantial heterogeneity in Wnt signaling both before and after the CHIR pulse (**Figure S1C**), consistent with a recent study^16^. In contrast, maintaining cells in 2i+LIF media until immediately before seeding (**Figure 1C**) produced gastruloids in which Wnt pathway activity indeed progressed from a uniform state at the end of the CHIR pulse to a single posterior pole of Wnt activity over time (**Figure 1D**). We proceeded with our modified protocol for all subsequent experiments to maintain tight control over the initial conditions of Wnt signaling while still supporting robust gastruloid development.

Wnt biosensor measurements over time revealed a progression through uniform, bimodal, and polarized pathway activity states. At 48 h post-seeding, prior to the CHIR pulse, Wnt activity was low in all cells but shifted to a uniformly-high state by 72 h at the end of the CHIR pulse (**Figure 1D**, left). By 96 h, we observed fragmentation of the Wnt activity pattern into locally ordered domains of active and inactive signaling, but without global axial polarization (**Figure 1D**, middle). Finally, by 108 h post-seeding, gastruloids exhibited a single coherent domain of Wnt activity that marked the elongating posterior. This sharply-defined domain of posterior Wnt activity persisted through subsequent time points, occasionally showing signs of further fragmentation with the emergence of new Wnt-inactive patches (**Figure 1D**, right). We separately characterized single-cell Wnt activity levels by applying flow cytometry suspensions of dissociated gastruloid cells (**Figure 1E-F; Figure S1D**). Consistent with our imaging data, these flow cytometric measurements also revealed an initial homogeneous low state at 48 h post-seeding, followed by a uniformly high state at 72 h, finally shifting to a bimodal distribution with Wnt-high and Wnt-low subpopulations at subsequent time points.

To pinpoint the timing of the transitions to the heterogeneous and polarized states, we prepared gastruloids expressing our instantaneous Wnt biosensor at 6 h time intervals between 72 h and 144 h post-seeding (**Figure 1G; Figure S1E**; n = 76 gastruloids) and imaged the spatial distribution of Wnt activity. We quantified polarization by measuring the center of mass of Wnt fluorescence compared to the ubiquitously expressed DsRed marker, and quantified heterogeneity using the standard deviation of normalized pixel intensities for each gastruloid (see **Figure S1F**; see **Methods**). These analyses revealed a sharp increase in Wnt heterogeneity at 90 h post-seeding (**Figure 1G**; red curve), consistent with our flow cytometry results (**Figure 1E**). In contrast, Wnt polarization to a posterior domain was delayed by an additional 12 h, only increasing at 108 h post-seeding (**Figure 1G**; blue curve). In summary, a live-cell signaling biosensor revealed that Wnt pathway activity is highly dynamic and can self-organize into a polarized pattern in response to a uniform cue (the CHIR pulse) by evolving through multiple intermediate states: (1) global low and high Wnt states from 48-90 h post-seeding, (2) heterogeneous, “patchy” Wnt activity without overall polarization from 90-108 h, and finally (3) posterior-polarized Wnt activity coinciding with the emergence of morphological polarization from 108 h onward. Notably, revealing this progression of patterns depends on a gastruloid preparation protocol that minimizes early Wnt heterogeneity^16^ and enables all cells to initially respond to the synchronizing CHIR pulse.

### Recording signaling histories with recombinase circuits

We observe that gastruloid Wnt activity undergoes a complex evolution from a homogenous state to a single posterior pole. Might the sequential spatial patterns of Wnt activity carry functional significance? Early cell-to-cell differences in Wnt activity may simply reflect transient fluctuations prior to a later decision timepoint, or they may already carry information about cells’ future position and fate. To test the hypothesis that early Wnt heterogeneity might already predict cell fates, we devised a general experimental strategy to (1) label cells based on their signaling state at precisely defined times, (2) continue to grow gastruloids until some final time of interest and (3) assess the final spatial position of these labeled cells in the end-stage gastruloid. Beyond mapping Wnt-associated cell positions, we reasoned that such a “signal-recording” system could be useful for tracing the fates of signaling-active cells for a broad range of pathways and biological contexts.

We adopted a generalizable strategy of gating the expression of a site-specific recombinase (e.g., Cre recombinase) to permanently label cells based on their signaling histories^25, 26^. To restrict labeling to a defined time window, we built a synthetic circuit where recombinase activity is controlled by an AND gate sensitive to two inputs: signaling pathway activity and an experimentalist-controlled small molecule, doxycycline. In principle, delivering the small molecule during a brief, well-defined time window would thus label only those cells currently in an active signaling state for analysis at later time points (**Figure 2A**). Similar approaches have been used *in vivo* to label signaling-active neurons during experimentally-defined time windows (e.g., the *Fos-tTA* system), but with temporal precision typically measured in days rather than hours^27^.

**Figure 2:**
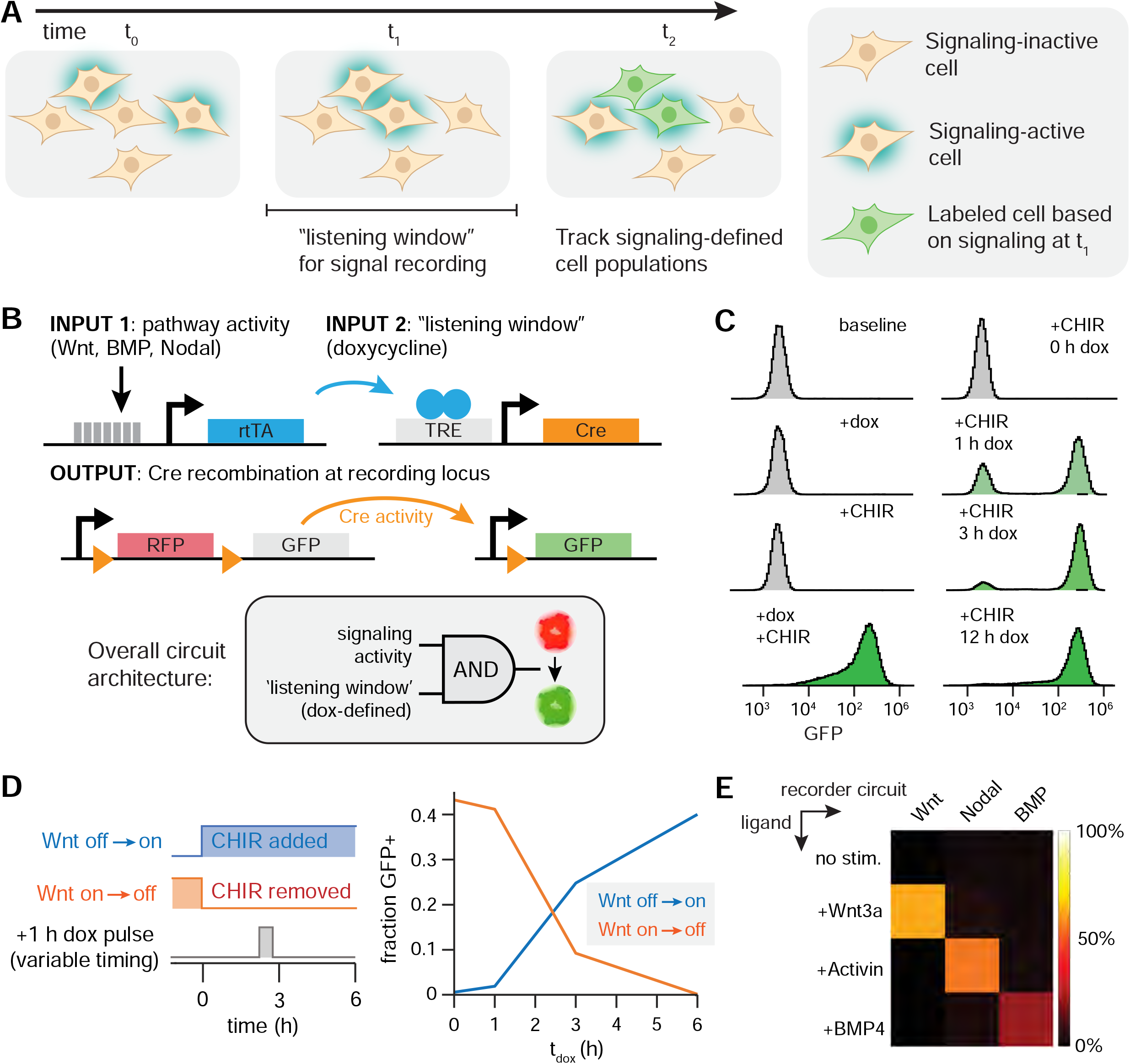
Recording morphogen signals with recombinase circuits. **(A)** Illustration of signal recording design criteria. An ideal recorder would irreversibly label a signaling-defined population within a temporal ‘listening window’ of interest so that this population can be followed over time. **(B)** Schematic of recording circuit. An upstream transcription factor (rtTA) requires both signaling activity and small-molecule addition to drive recombinase expression, which in turn irreversibly changes the fluorophore in a ‘recording’ locus. **(C)** Characterization of a clonal Wnt recording cell line fidelity. 24 h incubation with both Wnt-activating CHIR (3 µM) and listening window-defining doxycycline (2 µg/mL) achieves complete GFP labeling (left, bottom). Treatment with either CHIR or doxycycline alone has no detectable labeling (left, top). A 1 h recording window is sufficient to achieve efficient labeling (right). **(D)** Characterization of switching kinetics of Wnt recorders in response to media changes. A 1 h doxycycline treatment was applied at variable lag times (Δt = t_dox_ – t_0_) following a media change. Recorder performance approached steady-state media performance by Δt = 6 h. **(E)** Crosstalk assessment for 3 separate clonal lines recording Wnt, Activin/Nodal, and BMP pathway activity. All recording windows utilized 100 ng/mL doxycycline and 200 ng/mL morphogen concentration. Baseline conditions report labeling with only doxycycline in basal media (N2B27). Pseudocolors indicate relative proportion of cells labeled with GFP expression. Recording windows were 6 hours for Wnt and Nodal recorders and 3 h for the BMP recorder to account for differences in sensitivity.

We implemented our dual-input recombinase circuits in mouse embryonic stem cells using serial rounds of PiggyBac-mediated integration (**Figure 2B**)^28^. The top-most element in our circuit uses a signaling pathway-responsive ‘sentinel enhancer’^29^ (in our base case for the Wnt pathway, 7 TCF/LEF binding sites) to drive the expression of a destabilized reverse tetracycline dependent transactivator (rtTA-PEST). Only in the presence of the small molecule doxycycline will rtTA then drive expression of a TetO-driven Cre recombinase^30^. Finally, Cre recombinase can act on a “stoplight” recording cassette (P_EF1α_-LoxP-DsRed-LoxP-GFP)^31^, excising the DsRed transgene to permanently switch cells from red to green fluorescence. We first engineered a clonal mESC cell line expressing the P_TetO_ -Cre and stoplight elements to serve as a chassis for signaling-recorder circuits that respond to a variety of inputs, into which we integrated a signaling-responsive enhancer specific for either Wnt, Nodal or BMP signaling (see **Methods**). At each stage of cell line generation, mESC clonal colonies were selected for colony morphology and lack of leaky recombination-mediated GFP expression.

We characterized the fidelity of a “Wnt-Recorder” mESC clonal cell line by measuring the efficiency of labeling in response to both inputs. We treated 2D cultures of mESC with different combinations of doxycycline (dox; 2 μg/mL) or CHIR (3 μM) for 24 h, washed and returned to basal media, and then waited an additional 24 h to equilibrate recombination-mediated shifts in fluorescent protein expression prior to measuring GFP fluorescence by flow cytometry. We found that the circuit exhibited very low leakiness: no detectable GFP labeling was observed when Wnt-Recorder cells were incubated for 24 h in either dox or CHIR alone (**Figure 2C**), and background GFP levels remained consistently low (<0.1%) over at least 15 passages of subculture in Wnt-activating 2i+LIF media but in the absence of doxycycline. Conversely, combined stimulation with dox and CHIR produced near-complete GFP labeling of the cell population (**Figure 2C**, left). We further characterized the sensitivity of the circuit to each input (dox or Wnt signaling) in the presence of a constant cue from the other input (**Figure S2A-B**), revealing that labeling could be detected for doxycycline concentrations as low as 200 ng/mL and for Wnt3A concentrations as low as 50 ng/mL.

For high-fidelity signal recording it is important that a brief incubation with doxycycline is sufficient to label cells based on their current pathway activity. To measure the minimum recording window required for efficient labeling, we first varied the duration of doxycycline labeling in the presence of a constant CHIR stimulus (**Figure 2C**, right). We found that the circuit exhibited exceptional temporal sensitivity – even a 1 h incubation window was sufficient to label the majority (68%) of cells, with near-complete labeling after a 3 h dox pulse. As a more stringent test, we performed step-up and step-down experiments to measure recording kinetics after an acute change in Wnt activity. We incubated Wnt-Recorder mESCs with a 1 h pulse of doxycycline at various time points after either CHIR stimulation or washout, and measured GFP labeling after 24 h (**Figure 2D**). We generally observed a delay of 3 h before the doxycycline pulse produce labeling results that reflected the change in Wnt activity, likely reflecting the time required for synthesis or degradation of the Wnt-induced rtTA transcription factor. By 6 h following the media change, labeling fractions approached steady state values for low-Wnt or CHIR-stimulated conditions. We thus conclude that Wnt-Recorder cells can faithfully resolve signaling dynamics to within a 6 h time window.

Recombinase recording circuits are modular in design: exchange of the upstream most sentinel enhancer can target recording to different morphogen signals of interest. We further developed clonal mESC lines to record Activin/Nodal signaling using the AR8 sentinel enhancer and BMP activity using the IBRE4 sentinel enhancer^29^, terming the resulting mESC clones Nodal-Recorder and BMP-Recorder cells, respectively. All three Recorder cell lines (Wnt, Nodal, and BMP) could be used to faithfully record ligand-dependent signaling from the appropriate pathway with minimal crosstalk from the other two pathways (**Figure 2E; Figure S2C**). For recording windows of 6 h or longer, the BMP-Recorder line exhibited some recording in the absence of exogenous BMP4 (**Figure S2D**) but this basal activity was suppressed by treatment with the BMP inhibitor LDN-193189 and thus may accurately reflect autocrine BMP signaling within the culture (**Figure S2D**). In sum, the synthetic gene circuits described here enable the experimentalist to label mESCs based on their instantaneous Wnt, Nodal or BMP signaling activity with high fidelity and fine temporal resolution.

### Tracing the emergence of fate information in Wnt signaling

We next applied our Wnt-Recorder system to measure whether early signaling differences in this pathway are predictive of cells’ future axial position. We envisioned a simple experimental protocol: establish gastruloids from Wnt-Recorder cells, apply a 90 min pulse of doxycycline at a time t_dox_ to label cells based on their immediate history of Wnt activity, and finally fix and image the elongated gastruloids at a final time t_f_ (**Figure 3A**). If cells’ early Wnt signaling state is instructive of their final position, then recording Wnt-active cells and tracing their fates should reveal an asymmetric distribution of GFP-labeled cells along the gastruloid’s final A-P axis. We hypothesized that signal recorders could thereby measure ‘fate information’ encoded in cells’ signaling states; that is, information that does not encode a cell’s current position, but rather predicts where it will end up in the final polarized gastruloid.

**Figure 3:**
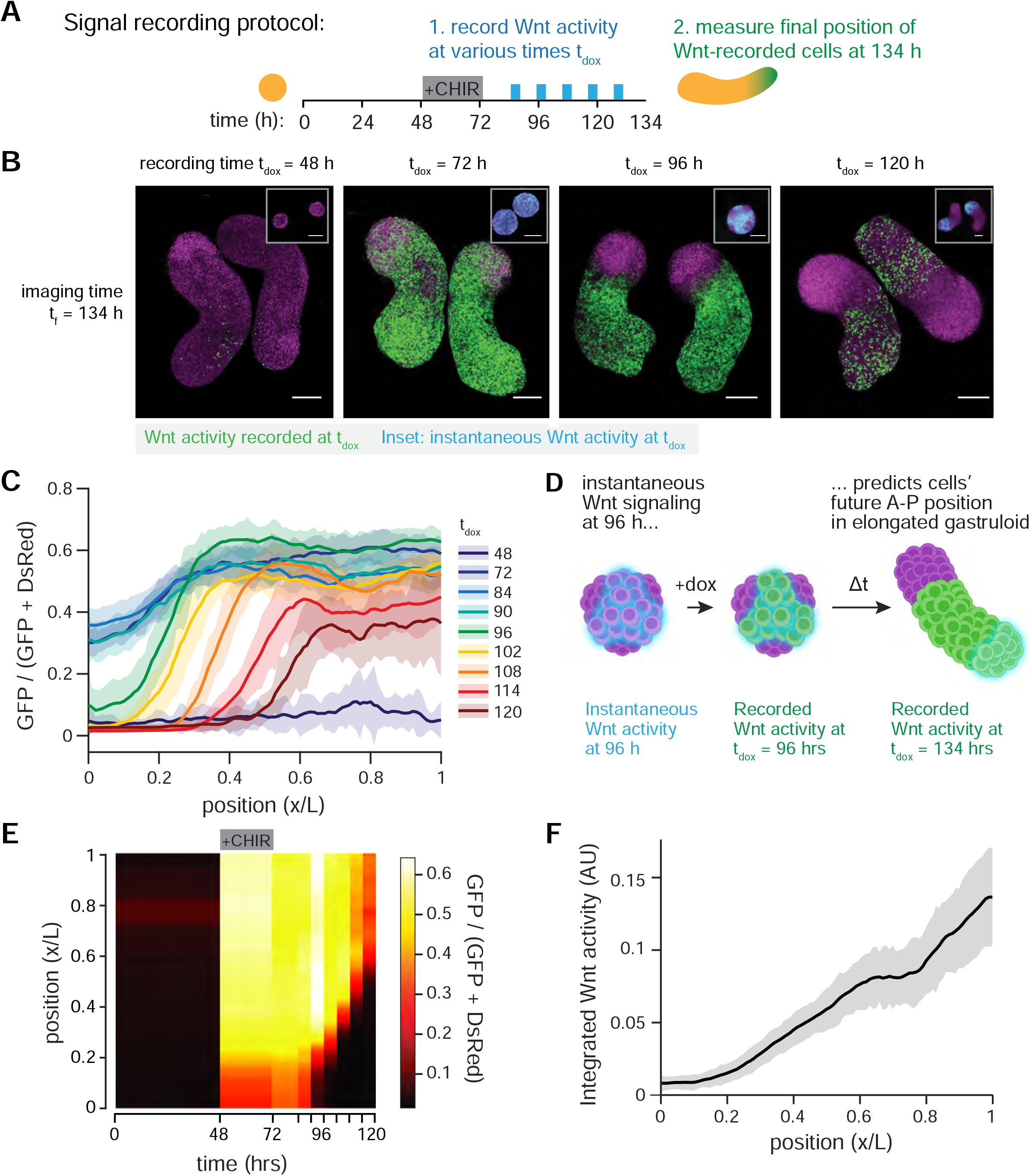
Mapping fate information encoded in Wnt signaling histories. **(A)** Schematic of experimental design. Wnt histories are recorded by varying the onset time t_dox_ of a doxycycline-defined listening window (90 minutes, 200 ng/mL) during gastruloid morphogenesis. **(B)** Representative final images (t_F_ = 134 h) of gastruloids in which Wnt activity was recorded at different timepoints. Insets (top-right) show the corresponding pattern of instantaneous Wnt activity (**Figure. 1D**) during the queried listening window (scale bar = 200 um). **(C)** Quantification of anterior-posterior (A-P) axial patterns of Wnt recorder labeling at different timepoints throughout gastruloid morphogenesis at 6 h temporal resolution (n = 119 gastruloids measured). Shaded regions indicate standard deviation as a function of A-P position. **(D)** Illustration of patterns corresponding to the ‘symmetry-breaking window’ at t_dox_ = 96 h, the first timepoint at which Wnt recording predicts a clear anterior-posterior separation of cell fates. The measurement of fate information within ‘patchy’ patterns of Wnt activity suggests cellular rearrangements contribute to gastruloid polarization. **(E)** Kymograph of the data presented in Figure 1C to visualize different Wnt dynamics corresponding to different spatial positions. Following the emergence of an anterior domain at t_dox_ = 96 h, Wnt signaling becomes progressively more restricted to the posterior domain. **(F)** Integrated Wnt signaling activity between t = 96 and t = 134 h shows a linear ‘temporal gradient’ associated with A-P fate.

We grew cohorts of Wnt-Recorder gastruloids and chose 9 time points between 48-120 h to apply doxycycline pulses for signal-recording. Recording timepoints were spaced by a minimum of 6 h, matching the sensitivity window of the Wnt-Recorder circuit (**Figure 2D**). All gastruloids were grown to a final fixation timepoint t_f_ between 134 and 144 h, and then imaged to observe the morphogenic fates of signal-recorded cell populations (**Figure 3B**). Applying the doxycycline pulse at 48 h resulted in gastruloids exhibiting only sparse and sporadic GFP labeling (**Figure 3B**, t_dox_=48 h), consistent with our observation of uniform, low Wnt activity at 48 h post-seeding (**Figure 3B**, inset above t_dox_=48 h). Conversely, a dox pulse delivered at 72 h produced GFP labeling throughout the entirety of the gastruloid, consistent with the high level of Wnt pathway activity expected in all cells immediately following CHIR treatment. We observed the emergence of axially-organized labeling patterns by t_dox_=96 h, where GFP-labeled cells were evenly distributed across much of the gastruloid but were entirely excluded from the anterior domain (**Figure 3B**, t_dox_=96 h). At progressively later timepoints, cells harboring Wnt signaling were increasingly restricted to the posterior of the gastruloid. We quantified data for all labeling times by measuring normalized GFP as a proportion of the total normalized intensity of the Stoplight system (GFP + DsRed) along the major axis of each gastruloid (**Figure 3C; Figure S3A-D**; see **Methods**). This analysis confirmed our raw imaging data (**Figure 3B**) that GFP labeling transitioned from a uniformly-low state for t_dox_=48 h to a high state at t_dox_=72-90 h, followed by progressive restriction to a posterior domain from t_dox_=96 h onward. We also noted a slight trend in the GFP/(GFP + DsRed) ratio at the anterior-most 10% of the gastruloid (**Figure 3C**; 72-90 h curves) that was present even at the earliest labeling time points when Wnt signaling was still uniform (**Figure 1E**). This trend may reflect intrinsic differences at the anterior pole (e.g., high cell density or autofluorescence) that affect the relative fluorescence measurements in the GFP and DsRed channels.

Our data indicate a stark transition between 90-96 h post-seeding where Wnt signaling first becomes associated with cells’ future position along the anterior-posterior axis (**Figure 3D**). Importantly, Wnt heterogeneity is not yet spatially organized along an anterior-posterior axis at this time, but rather Wnt-high and Wnt-low cells coexist in multiple patches across a single gastruloid without a clear axial bias (**Figure 1D-G**). Put differently, by 96 h, cells with high Wnt activity are already destined to be excluded from the gastruloid’s anterior-most domain, even though these cells are not yet spatially organized along an A-P axis.

In addition to assessing the future position of Wnt-active cells at a single labeling time, our doxycycline-labeling experiments can also be re-cast as measuring the dynamic Wnt signaling history for cells at each final position in the gastruloid. This can be visualized by considering horizontal slices through a kymograph of our Wnt labeling data, representing the Wnt activity over time experienced by cells at each final position (**Figure 3E**). For example, a high proportion of cells at the gastruloid’s posterior-most positions (i.e., x/L = 1) were Wnt-active from 72-120 h, whereas cells at the midpoint (x/L = 0.5) saw high Wnt activity only from 72-108 h. We thus set out to ask whether each position in the final gastruloid might be well-defined by its dynamic Wnt signaling history. To address this question, we quantified the total integrated Wnt signaling observed at each spatial position:

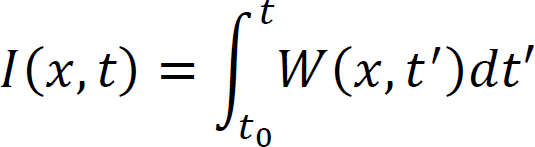

where *W*(*x, t*’) is a normalized position-dependent distribution of Wnt activity measured at time *t*’ Computing this integral from t_0_ = 96 h through t = 134 h yielded a roughly linear gradient (**Figure 3F**). These results reveal a ‘temporal gradient’ of prior Wnt activity at each A-P position in the final gastruloid: cells at each axial position can be distinguished by the total duration of Wnt signaling they have experienced. Because instantaneous Wnt activity in the elongating gastruloid (> 108 h post-seeding) is restricted to the posterior pole (**Figure 1D; Figure S3D**), it is likely that this temporal gradient reflects the dynamics with which newly born cells exit the posterior-most Wnt domain during gastruloid elongation, turning off this signaling pathway as they differentiate and contribute to tissues at various axial positions.

Our signal-recorder observations are already sufficient to begin constraining potential mechanisms of gastruloid symmetry breaking. First, we may infer that gastruloid symmetry breaking must include at least some contribution from cell reorganization, since Wnt-high cells that are present throughout the gastruloid at 96 h must subsequently sort away from the future anterior. Such a mechanism would argue against a purely biochemical Turing-like reaction-diffusion system, operating on a fixed field of cells, in setting up a single polarized anterior or posterior domain. Second, our data indicates that a subpopulation of cells retains persistent, high Wnt activity after removal of the CHIR pulse, and that this Wnt-high state persists long enough to transition from patches throughout the gastruloid to a single posterior domain. The long-term maintenance of a Wnt-high state might indicate the presence of local positive feedback acting within the Wnt-high subpopulation^21^. More broadly, our results demonstrate the utility of signaling-recorder gene circuits for defining when signaling pathway activity is first informative of symmetry-breaking polarization.

### Early Wnt activity is associated with distinct cell fate trajectories

We have seen that cell-to-cell differences in Wnt signaling at 96 h post-seeding are predictive of cells’ future position in the elongated gastruloid; are they also associated with the adoption of specific cell fates? We reasoned that combining our signal-recorder cell lines with single-cell RNA sequencing (scRNAseq) could address this question: cells can be GFP-labeled based on their instantaneous signaling state, and then harvested at any future time of interest to assess their transcriptomic state. We thus repeated the 96 h labeling of Wnt-Recorder gastruloids that also expressed the instantaneous destabilized iRFP Wnt reporter, then dissociated and sorted the GFP+ and GFP-subpopulations at 120 h for scRNAseq (**Figure 4A**; see **Methods**). Imaging experiments confirmed that by 120 h post-seeding, gastruloids exhibited high instantaneous Wnt signaling at the posterior pole as well as GFP labeling that was excluded from the anterior domain (**Figure 4B**).

**Figure 4:**
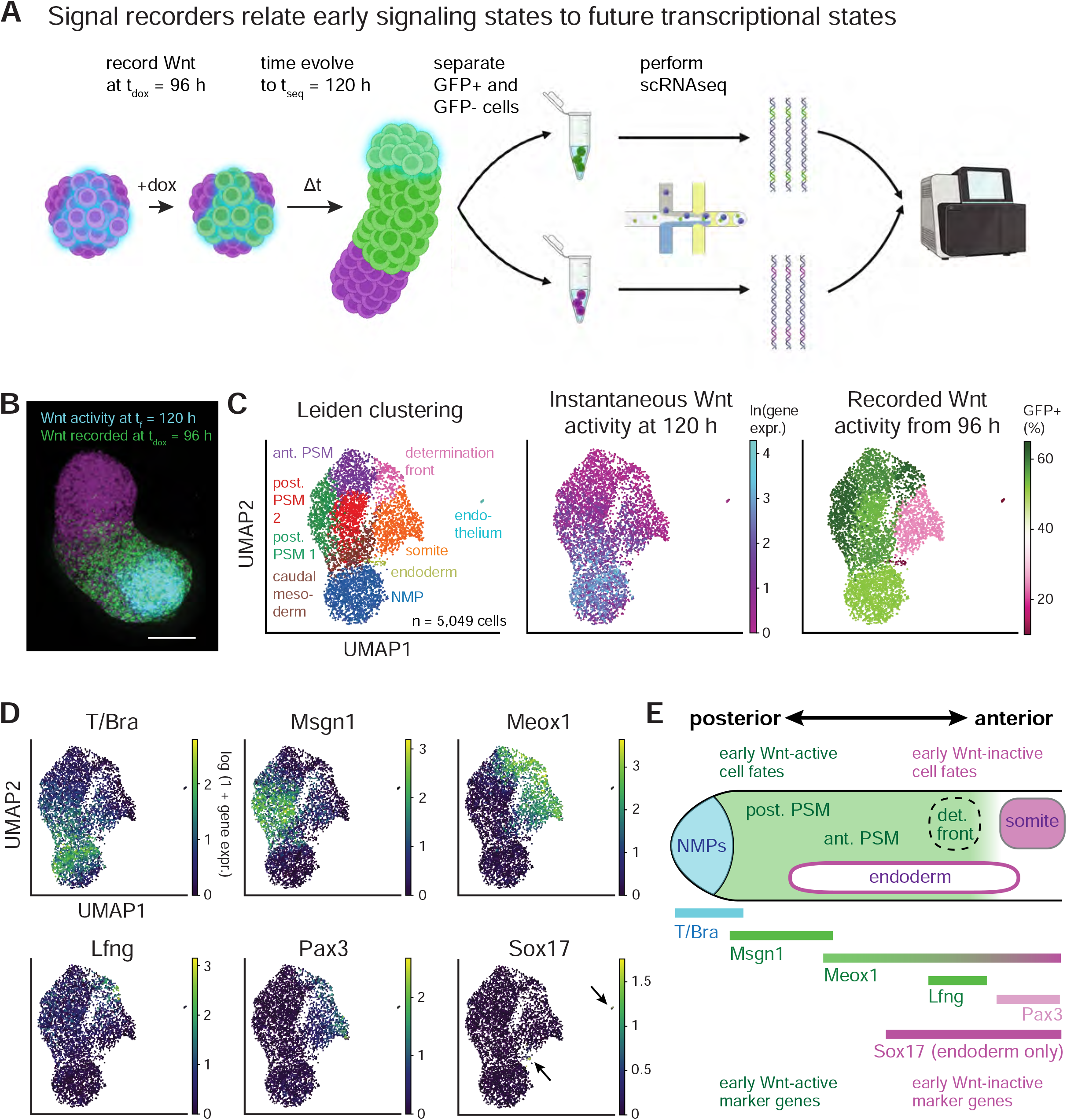
Mapping signaling histories in transcriptional space. **(A)** Schematic of experimental design. Wnt signaling states are recorded in a 90 minute window initiated at t_dox_, and gastruloids are then grown to a later time t_seq_. Gastruloids are then dissociated into single cell suspensions, and separated into GFP positive and negative signaling populations via fluorescence activated cell sorting (FACS). Sorted populations are then loaded into separate lanes of a Chromium controller (10× Genomics) to prepare for single-cell RNA sequencing (see Methods). Separate gene expression libraries are then prepared with distinct library indices to disambiguate signal recording conditions in pooled sequencing. **(B)** Representative image of a gastruloid at t_seq_ = 120 h, labeled according to signaling domains (scale bar = 200 µm). **(C)** Annotating cell type clusters with signaling information. Left: cell types identified by Leiden clustering within t_seq_ = 120 h gastruloids. Middle: final Wnt activity distribution across cells as measured by P_TCF/LEF_-rtTA expression. Final Wnt activity is concentrated within the neuromesodermal progenitor (NMP) cluster. Right: relative composition of Leiden clusters according to Wnt activity recorded at t_dox_ = 96 h. Wnt activity is broadly distributed throughout most clusters, but excluded substantially from the somite, endoderm, and endothelial clusters (see also **Methods**). **(D)** Single-cell expression levels of reference genes associated with gastrulation and axial morphogenesis. **(E)** Illustration of reference gene organization across the anterior-posterior axis during axial elongation and somitogenesis. The organization of reference gene expression (Figure 1D) aligns with inferred histories of Wnt activity (Figure 1C), suggesting that Wnt signaling histories predict transcriptionally defined cell fates along a spatially defined anterior-posterior axis.

The combined scRNAseq dataset from GFP+ and GFP-cells revealed 9 cell clusters (**Figure 4C**, left panel) that could be assigned to distinct embryonic cell types using annotations from a reference scRNAseq atlas mapping mouse development between embryonic day 6.5 and 8.5 (**Figure S4A**; see **Methods**)^32^. The cell types which we observed were largely consistent with those previously reported in similarly staged gastruloids^33^. These fates were also associated with a progression along the posterior-to-anterior axis, from the posterior-most neuromesodermal progenitors (NMPs) to caudal and presomitic mesoderm (PSM), followed by anterior mesoderm and cell types associated with somitogenesis (i.e., ‘determination front’ and ‘somite’ populations). We also observed compact clusters of endodermal and endothelial cells whose expected position along the A-P axis is less well-defined. Overall, the cell types annotations were broadly consistent with the expression profiles of known marker genes (**Figure 4D; Figure S4B**), with T/Bra marking NMPs and posterior PSM, Msgn1 marking posterior PSM, Meox1, Lfng, and Pax3 marking cells undergoing somitogenesis, and Sox17 expressed exclusively in the endodermal and endothelial clusters. Cell type annotations were also consistent with the expected localized expression of signaling-associated genes (**Figure S4C**), with high expression of Wnt and FGF ligands (Wnt3a, Wnt5b, Fgf8, and Fgf17) in posterior-associated cell clusters and retinoic acid synthesis (Aldh1a2) in anterior-associated clusters, matching the opposing gradients of Wnt/FGF and retinoic acid signaling along the A-P axis^34, 35^.

Our scRNAseq analysis further revealed that both instantaneous and recorded Wnt signaling map to specific cell types in the 120 h old gastruloid. We observed high levels of instantaneous Wnt activity (measured by TCF/LEF-driven rtTA transcription) in the NMP and caudal mesoderm subpopulations (**Figure 4C**, middle panel), consistent with the spatial distribution of instantaneous Wnt signaling at the posterior pole (**Figure 4B**) where NMPs are located^33^. While both GFP+ and GFP-cells were found in similar proportions throughout each cluster (**Figure S4D-E**), we found that GFP-expressing cells were largely excluded from three clusters: the anterior-most ‘somite’ cluster^33^ and the ‘endoderm’ and ‘endothelium’ clusters whose axial localization patterns are less well-defined (**Figure 4C**, right). Taken together, these data reveal that cell-to-cell heterogeneity in Wnt signaling at 96 h is indeed associated with distinct subsequent differentiation trajectories (**Figure 4E**). Cells with high Wnt activity at 96 h contribute exclusively to NMP and posterior mesodermal fates, whereas low Wnt activity at 96 h is already predictive of anterior mesoderm and endodermal/endothelial lineages. More broadly, these data demonstrate that signal-recorder circuits can be combined with standard scRNAseq pipelines to annotate maps of cell fates with information about their current and prior signaling histories.

### Symmetry breaking and polarization involves differential adhesion

Our prior experiments reveal a critical time window from 90-96 h when heterogeneity in Wnt signaling becomes predictive of cells’ future axial positions and transcriptional fates in the elongated gastruloid. This time window corresponds to a period during which Wnt signaling exhibits local order in patchy domains but has not yet polarized to the posterior of the gastruloid (**Figure 5A**). We next investigated how these Wnt-high and Wnt-low subpopulations differ during the critical window to better understand how their subsequent polarization might occur.

**Figure 5:**
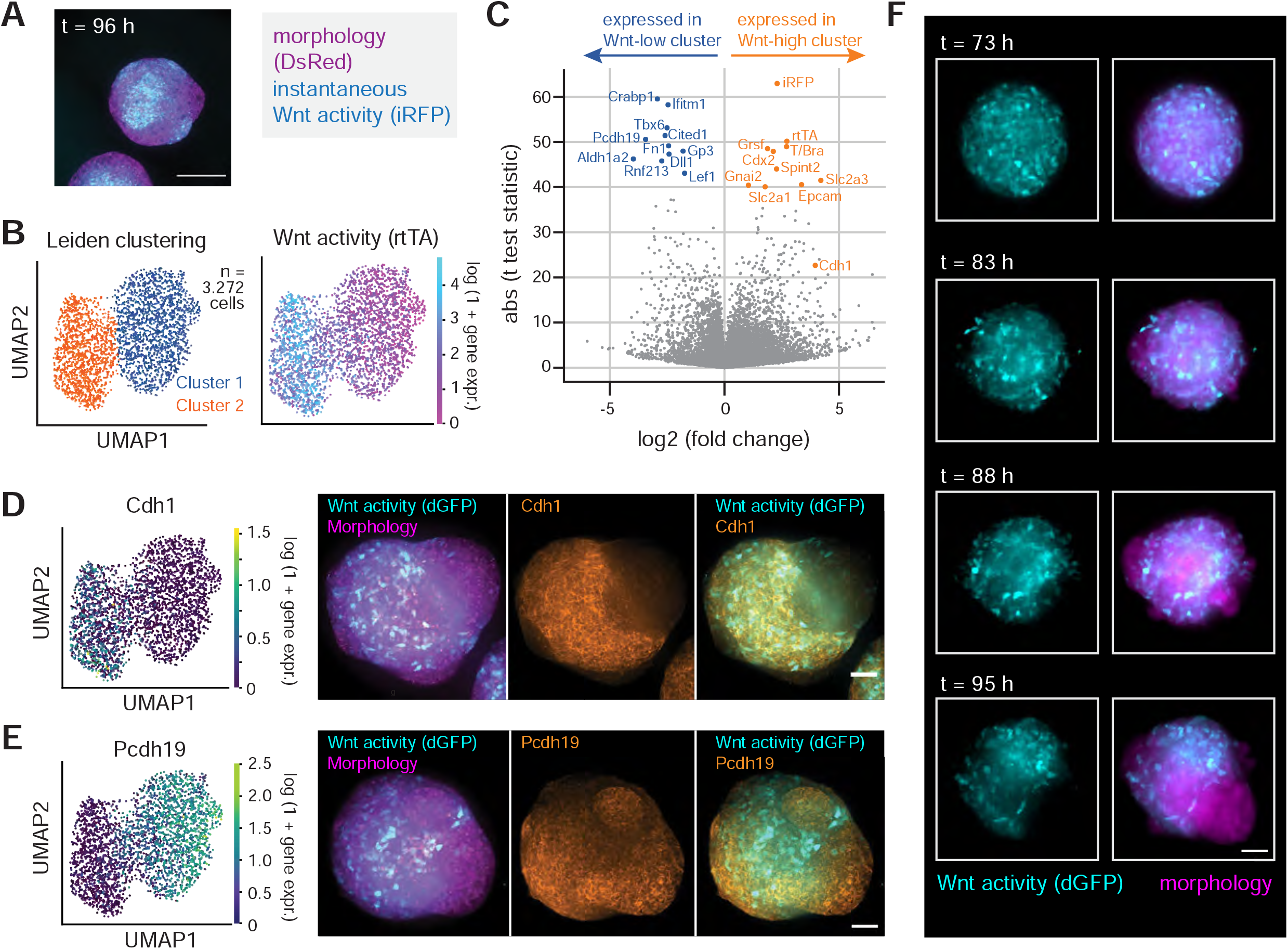
Wnt signaling during symmetry breaking is associated with differential transcriptional and mechanical cell states. **(A)** Wnt activity at t = 96 h is organized into ‘patchy’ patterns of local correlated domains lacking global polarization. **(B)** scRNAseq analysis of gastruloids collected at t_seq_ = 96 h. Left: Leiden clustering identified 2 cell types. Right: Wnt activity (as measured by P_TCF/LEF_-rtTA expression) was highly concentrated within cluster 2 (‘Wnt active’) and excluded from cluster 1 (‘Wnt inactive’). **(C)** Volcano plot reveals genes which are differentially expressed between the two Leiden clusters. **(D-E)** Validation of differential cadherin expression between cell clusters by immunofluorescence. **(D)** The Wnt-active cell type is characterized by exclusive expression of E-Cadherin/Cdh1. **(E)** In contrast, the Wnt-inactive cell type is enriched for Protocadherin-19/Pcdh19, an alternative homotypic adhesion marker. **(F)** Still images from **Movie S1** illustrating dynamics of cellular rearrangements during Wnt polarization. These rearrangements sort cells from patchy expression domains into a globally polarized signaling axis (Figure 3D).

To fully characterize the transcriptional state of the Wnt-high and Wnt-low populations, we performed scRNAseq on 96 h old Wnt-Recorder gastruloids also expressing our instantaneous Wnt biosensor (7×TCF/LEF-driven iRFP-PEST). We performed Leiden clustering on the resulting 3,272 single-cell transcriptomes, which identified two populations of cells which differentially express many genes previously identified as marker genes in a recent mouse developmental atlas(**Figure 5B-C; Figure S5A**)^32^. Reads corresponding to our Wnt transcriptional biosensor were confined to just a single cluster, suggesting that the Wnt-high and Wnt-low signaling domains evident by microscopy also represent the primary distinguishable cell populations in this transcriptomic dataset. These two clusters were also associated with differential expression of a broader range of signaling-associated genes (**Figure S5B**). For example, the Wnt-high cluster expressed the Wnt ligands Wnt3a and Wnt5b, as well as BMP and FGF ligands, consistent with autocrine positive feedback locally maintaining Wnt activity in this subpopulation after CHIR washout at 72 h. Conversely, the Wnt-inactive population expressed a distinct set of signaling-associated genes, including the Wnt inhibitor FrzB and the retinoic acid biosynthesis enzyme Aldh1a2 (**Figure S5D**), which plays an opposing role to FGF signaling during somitogenesis^36^ and might reflect a nascent somitogenic program is already active by this stage of gastruloid development.

We also observed differential expression of cell adhesion-associated genes between Wnt-high and Wnt-low clusters. E-Cadherin/Cdh1 and EpCAM were expressed in the Wnt-high cluster, whereas protocadherins 8 and 19 (Pcdh8; Pcdh19) were predominantly expressed in Wnt-low cells (**Figure 5C-D; Figure S5C**). Not all adhesion receptors were differentially expressed; for example, N-Cadherin/Cdh2 expression was observed uniformly across all cells (**Figure S5C**). We further validated that the differential gene expression we observed also extended to protein distributions by immunostaining for a subset of our hits, including Cdh1, Pcdh19, and Aldh1a2 (**Figure 5D; Figure S6**). In each case, we observed a tight correlation between the spatial domains of our instantaneous Wnt reporter and the candidate gene product. Overall, these data demonstrate that, by 96 h, cells occupying Wnt-high and Wnt-low patches already exhibit broad differences in signaling state and gene expression, consistent with their adoption of distinct future fates and positions.

The observation of differential expression of adhesion-associated genes lent further support to a model where the Wnt-high and Wnt-low subpopulations might sort out into distinct posterior and anterior domains, respectively^37, 38^. To obtain direct evidence of cell sorting, we performed live imaging of gastruloids grown from a clonal mESC line expressing a destabilized GFP reporter of Wnt activity and constitutive mCherry expression during the period at which symmetry breaking occurs (t = 72 to 96 h) (**Figure 5F; Videos S1 and S2**). We embedded gastruloids in 50% Matrigel to restrict their movement during live imaging^17, 18^. Wnt activity began broadly distributed throughout the gastruloid, with heterogeneous reporter expression levels in individual cells. Wnt-active cells were highly motile and, in the gastruloids where symmetry breaking could be observed (**Video S2**, arrows), gradually coalesced into clusters that merged into a single ring of cells, eventually extruding a broad domain of Wnt-inactive cells on one side of the gastruloid. Because Matrigel embedding may itself influence gastruloid development (e.g., by perturbing mechanical boundary conditions), we further scrutinized the intermediate patterns between domain emergence and polarization in fixed samples that were not embedded in Matrigel (**Figure S5E**). We observed a similar evolution of Wnt activity in these fixed samples, progressing from local patches to a ring, followed by resolution into a singular pole of Wnt activity. Overall, our live-imaging data provides direct evidence for sorting of Wnt-positive and Wnt-negative cells during gastruloid polarization, confirming the sequence of events that can be inferred from our signaling-recorder measurements.

In sum, single-cell gene expression analysis during the onset of symmetry breaking demonstrates that early heterogeneity in Wnt signaling is also associated with major differences in cells’ transcriptional states. Live imaging reveals that a cell sorting process rearranges cells harboring these different Wnt signaling states into progressively more consolidated domains, eventually yielding a polarized pattern of activity which defines a single posterior pole. Throughout this process, local patches of Wnt-high cells may be maintained through local secretion of Wnt3a and Wnt5b ligands, helping to maintain this subpopulation after removal of the CHIR pulse^21^. The observation that the two transcriptionally defined cell types express differential cell adhesion molecules and consolidate into distinct domains suggests that mechanical interactions could contribute to the cell rearrangements underlying gastruloid polarization and patterning.

### Spontaneous TGFβ signaling precedes and predicts Wnt symmetry breaking

Our data indicate that gastruloid polarization evolves from a uniform Wnt-high state to a bimodal population of Wnt-low and Wnt-high cells, followed by rearrangement of these cells into distinct anterior and posterior domains. The emergence of a bimodal distribution from uniformly Wnt-high initial state indicates that some cells maintain Wnt signaling longer than others following CHIR washout. We next sought to determine what cell state variables might contribute to this difference. Are there hidden variables in a cell’s state that might already predict Wnt persistence and thus A-P position even before CHIR addition at 48 h post-seeding?

We hypothesized that pre-existing heterogeneity in other developmental signaling pathway activity could bias cells to respond differently to CHIR stimulation. Indeed, Wnt signaling is only one component of a broader signaling network that organizes anterior-posterior axis specification during gastrulation *in vivo* (**Figure 6A**). The primitive streak is specified in the posterior domain of the epiblast not only by secretion of Wnt3 from the visceral endoderm, but also by secretion of BMP4 from the extraembryonic ectoderm. Within the nascent primitive streak, its anterior-most compartment (the ‘node’) is further influenced by Nodal signaling^39^. Both Nodal and BMP4 are TGFβ family ligands, and recent studies in human ES cell cultures indicate that TGFβ treatment can influence a cell’s response to a subsequent Wnt stimulus^40^. We therefore hypothesized that early cell-to-cell differences in Nodal and/or BMP signaling could serve as sources of heterogeneity to help break symmetry in response to Wnt stimulation.

**Figure 6:**
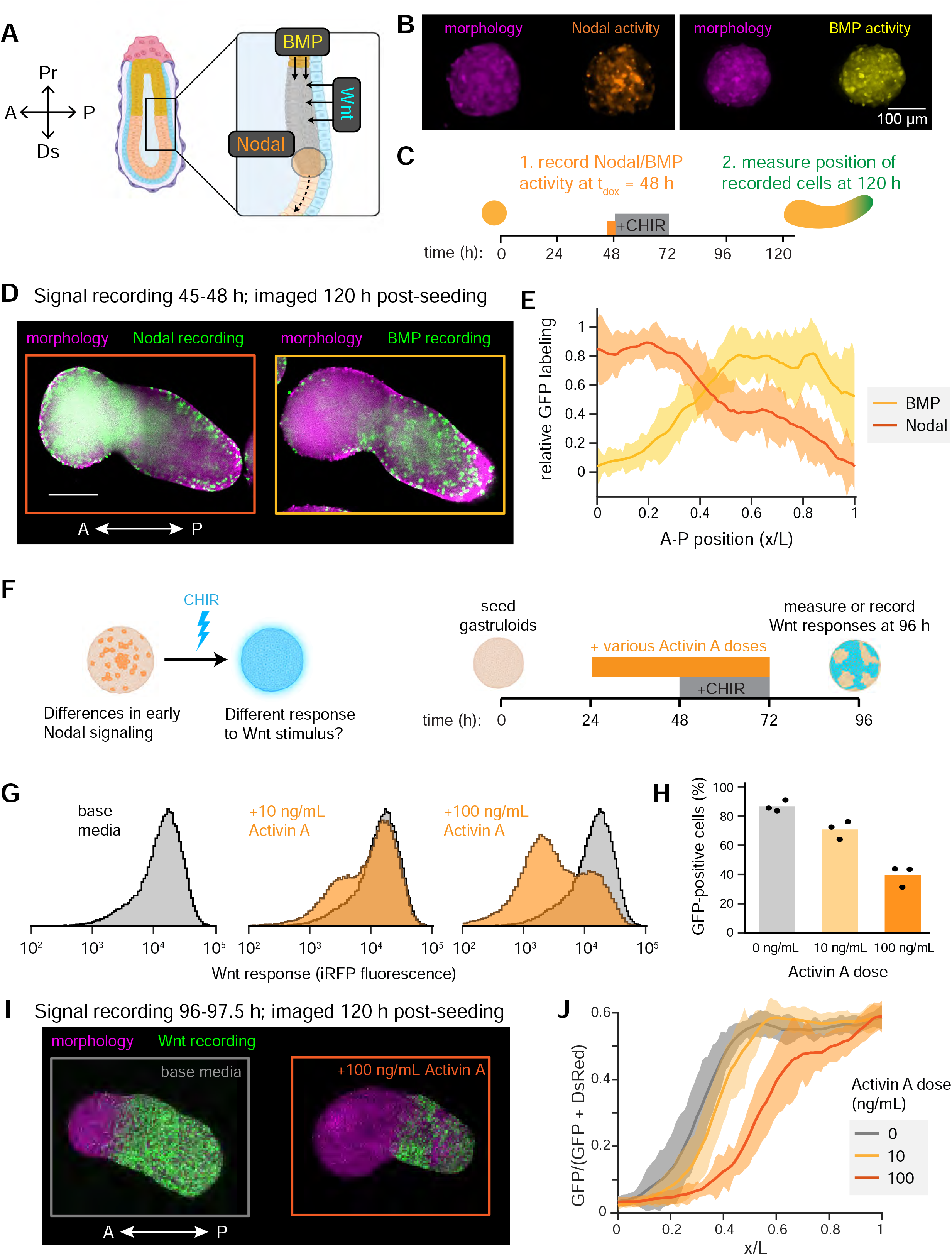
Early Nodal/BMP activity predicts and controls Wnt symmetry breaking. **(A)** In the mouse embryo, BMP and Wnt pathway signals interact in the posterior epiblast to initiate gastrulation. Nodal activity marks the anterior-most aspect of the resultant primitive streak (i.e., the ‘node’). **(B)** Gastruloids show spontaneous signaling activity in both the Nodal and BMP pathways at t = 48 h, before Wnt activity is detectable. **(C)** Signal recording experimental design. Early Nodal or BMP activity was recorded immediately prior to the stimulation of Wnt activity with CHIR (3 h recording window from t = 45-48 h; 200 ng/mL dox). **(D)** Representative images of final distribution (t_f_ = 120 h) of cells in which early Nodal/BMP activity was recorded, visualized through a medial optical section (scale bar = 200 µm). **(E)** Quantification of fate information recorded from early Nodal and BMP activity reveals that these predict a future anterior-posterior axis prior to the observation of Wnt activity (n = 22 gastruloids total). **(F)** Left: Illustration of a model in which differences in early Nodal activity influences differential future responses to Wnt stimulation, thereby predicting future cell fates. Right: experimental design to assess this model. **(G)** Single-cell Wnt activity levels at t = 96 h measured by flow cytometry. Activin A pretreatment drives more cells into the iRFP-negative (i.e. ‘Wnt-inactive’) population in a dose-dependent manner. **(H)** Quantification of the fraction of iRFP-positive (i.e. ‘Wnt-active’) cells across n = 3 replicates for each treatment condition (n = 30 pooled gastruloids per condition per replicate). **(I)** Representative images of gastruloids in which Wnt activity was recorded at t = 96-97.5 h and imaged at t = 120 h, with and without Activin A pretreatment. Scale bar = 200 µm. **(H)** Quantification of Wnt recording patterns demonstrates that Activin A pretreatment expands the Wnt-inactive anterior region (n = 42 gastruloids total).

To characterize Nodal and BMP signaling dynamics, we engineered clonal cell lines expressing instantaneous reporters of Wnt activity (P_TCF/LEF_-GFP-PEST) and an additional instantaneous reporter of either Nodal activity (P_AR8_-mCherry-PEST) or BMP activity (P_IBRE4_-mCherry-PEST) at distinct loci using separate piggyBac transformations. An additional constitutive marker (P_CMV_-TagBFP) marked overall gastruloid morphology. At t=48 h (i.e. just prior to CHIR application), we observed spontaneous activity in both pathways (**Figure 6B; Figure S6A-B**). Whereas BMP-active cells were dispersed throughout the gastruloid cross-section, Nodal activity appeared to be predominantly localized to a single domain. At 72 h post-seeding, immediately following CHIR stimulation, Nodal activity still showed a localized spatial pattern but with diminished amplitude (**Figure S6A**); conversely, BMP activity was elevated throughout the gastruloid at 72 h (**Figure S6B**). Finally, by 96 h post-seeding, both Nodal and BMP activity were reduced to uniform low levels (**Figure S6A-B**). These measurements demonstrate that although Wnt signaling evolves from an initially uniform state in our gastruloid protocol, the Nodal and BMP pathways also exhibit early signaling dynamics and cell-to-cell variability. Interestingly, this heterogeneity resolves to a uniform Nodal/BMP-off state by the time that Wnt signaling becomes spatially organized.

To understand whether the early heterogeneity in Nodal/BMP signaling might carry information about cells’ future A-P position, we prepared gastruloids from both Nodal-and BMP-Recorder cell lines, recorded signaling activity with a 3 h pulse of doxycycline from 45–48 h, and imaged the final recordings in elongated gastruloids at 120 h (**Figure 6C**). Mapping the positional distribution of GFP-labeled cells indeed revealed that the descendants of early Nodal-high and BMP-high cells indeed occupied specific positions along the A-P axis (**Figure 6D-E**). Cells labeled based on their Nodal activity were enriched in the anterior compartment of the final gastruloid; conversely, cells labeled with a BMP recording during the same window mapped onto the posterior domain. Remarkably, these opposing distributions of signaling-active cells from the two pathways suggest that gastruloids recapitulate the opposing Nodal/BMP patterning system found *in vivo*, despite lacking spatial pre-patterns of either signal from extraembryonic tissues. We performed additional Nodal/BMP labeling experiments at later times (**Figure S6C**), revealing that later BMP signaling continued to be associated with posterior signaling over time whereas Nodal signaling shifted to mark medial positions. In both cases, GFP labeling was lost as both pathways became inactive by 84-96 h.

Our data indicates that posterior positions in a final, elongated gastruloid are enriched for cells with high BMP signaling and low Nodal signaling at 48 h, as well as high cumulative Wnt signaling from 96 h onward. We next hypothesized that there might be a causal link between these sequential signaling states – that is, early differences in Nodal/BMP signaling might produce subsequent differences in Wnt signaling. To test this hypothesis, we established sequential stimulation protocols to first activate Nodal/BMP signaling and then monitor subsequent changes in Wnt pathway activity. We first grew gastruloids from Wnt-recorder cells that also expressed the instantaneous Wnt reporter, treated them with recombinant Activin A (a Nodal analog) or BMP4 from 24-72 h post-seeding, and measured their Wnt pathway transcriptional activity by flow cytometry at 72 and 96 h (**Figure 6F**). We found that cells’ initial Wnt response to CHIR stimulation was unaffected by Nodal/BMP pre-stimulation, with both treatment conditions still showing high, uniform Wnt responses at 72 h (**Figure S6E-F**). However, the duration of Wnt signaling following CHIR washout was sensitive to cells’ prior Nodal signaling state: Activin A pretreatment produced a consistent and dose-dependent change in Wnt signaling levels at 96 h, with a larger fraction of Wnt-off cells at increasing Activin A doses (**Figure 6G-H**). In contrast, BMP pretreatment exerted a more ambiguous effect. In gastruloids, BMP pretreatment partially reduced the magnitude of Wnt signaling in all cells but did not produce a bimodal response of distinct Wnt-on and Wnt-off subpopulations (**Figure S6F**). Subsequent experiments in cultured mESCs revealed that BMP could either potentiate or inhibit Wnt activity in a complex, dose- and time-dependent manner (**Figure S6G**), consistent with recent data from human ESCs^41^. Based on these data, we concluded that cells with high early Nodal signaling experience a more rapid loss in Wnt activity after CHIR washout. This ordered progression of cell signaling also matches the spontaneous activity in both pathways in gastruloids, with heterogeneous early Nodal activity followed by bimodal Wnt populations that sort to distinct anterior and posterior domains.

We reasoned that if Nodal activity decreased the probability of cells retaining a Wnt-high state, then gastruloids pretreated with increasing doses of Nodal should have fewer Wnt-high cells, resulting in a smaller posterior domain of Wnt activity. To test this prediction, we again pre-treated gastruloids with variable amounts of Activin A from 24-72 h, but this time we recorded Wnt activity at t_dox_ = 96 h and traced the fates of Wnt-labeled cells until t_f_ = 120 h. Gastruloids grown without Activin A pretreatment continued to form a small anterior domain from which Wnt recording at 96 h was excluded (**Figure 6I**, left), matching with our previous results (**Figure 3B-C; Figure 4B**). Consistent with our prediction, pretreatment with increasing doses of the Nodal analog Activin A altered these proportions to produce gastruloids with smaller Wnt-high posterior domains (**Figure 6I-J**). These data demonstrate a causal effect of Nodal pretreatment on the balance of anterior and posterior fates in the developing gastruloid.

Taken together, our experiments reveal an ordered sequence of signaling events that are associated with gastruloid symmetry breaking. Cell-to-cell differences in Nodal and BMP activity are already present in early mESC cell spheroids by 48 h post-seeding, prior to the CHIR pulse that initiates symmetry breaking and polarization through the Wnt pathway. These early signaling events go on to alter cells’ Wnt activity at 96 h, a time point where Nodal and BMP activity are no longer present, and alter the balance of anterior and posterior positional identity in the resulting gastruloid. In the vertebrate embryo, spatial pre-patterning of Nodal and BMP is used to define the coordinate system for gastrulation and axis formation. Our data reveals that gastruloids can use the same signaling network for polarization without an extraembryonic prepattern, suggesting that the same signaling network also possesses the capacity for self-organization of an anterior-posterior axis.

## Discussion

During embryonic development, a complex body plan emerges from comparatively simple initial conditions. Understanding how early cell signaling dynamics encode this complexity is a fundamental question in developmental biology. Programs of morphogenesis are frequently guided by spatially localized cues (e.g., from extra-embryonic tissues) which define patterned signaling domains. The case of the gastruloid offers a remarkable counter-example: mouse embryonic stem cells can self-organize a body axis *in vitro* even without any exogenous patterning cues. How do differences in cellular states break symmetry to define future differences in cells’ final positions or gene expression patterns?

Here we trace the signaling histories of mouse stem cells to explore how early heterogeneity in signaling pathway activity predicts future spatial patterning of the gastruloid body axis, an experimentally tractable *in vitro* model of developmental symmetry breaking. Our approach hinges on precise measurements of two kinds of information: (1) observing instantaneous patterns of signaling activity, and (2) making “recordings” of prior signaling activity at specific times to observe cells’ eventual position conditioned on their earlier activity states (i.e., ‘fate information’). We identify a set of gastruloid culture conditions under which we observe the onset of symmetry breaking in the Wnt signaling pathway, which progresses from an initial uniform state to intermixed domains of cells with either high or low Wnt activity. Synthetic Wnt-Recorder gene circuits reveal that this transition to bi-modality is accompanied by the emergence of fate information in Wnt signaling, such that cells’ Wnt signaling states at this time predict their future organization along the anterior-posterior axis. Only cells that have entered the Wnt-low state go on to form the gastruloid’s anterior domain and contribute to endoderm and endothelial fates, whereas Wnt-high cells occupy posterior positions and populate mesodermal and ectodermal lineages. We further show that the emergence of heterogeneity in Wnt signaling is influenced by prior heterogeneity in TGFβ signaling (e.g., Nodal and BMP pathway activity).

Our data suggests that an ordered sequence of signaling and cell rearrangement events drive gastruloid symmetry breaking (**Figure 7**). Pre-existing heterogeneity in spontaneous Nodal/BMP signaling is already present in spheroids of mouse embryonic stem cells, possibly due to stochastic gene expression or culture geometry (e.g. whether cells occupy exterior or interior positions in colonies). These pre-existing differences alter the duration of Wnt signaling following the application of a uniform CHIR pulse. The resulting Wnt-high and Wnt-low subpopulations express different adhesion molecules which may facilitate their spatial sorting into anterior and posterior domains, where they go on to assume distinct cell fates. Once the sorting process resolves in a single pole of Wnt activity, gastruloids can begin axial elongation through asymmetric expression of additional inductive cues such as FGF^17, 21^.

**Figure 7:**
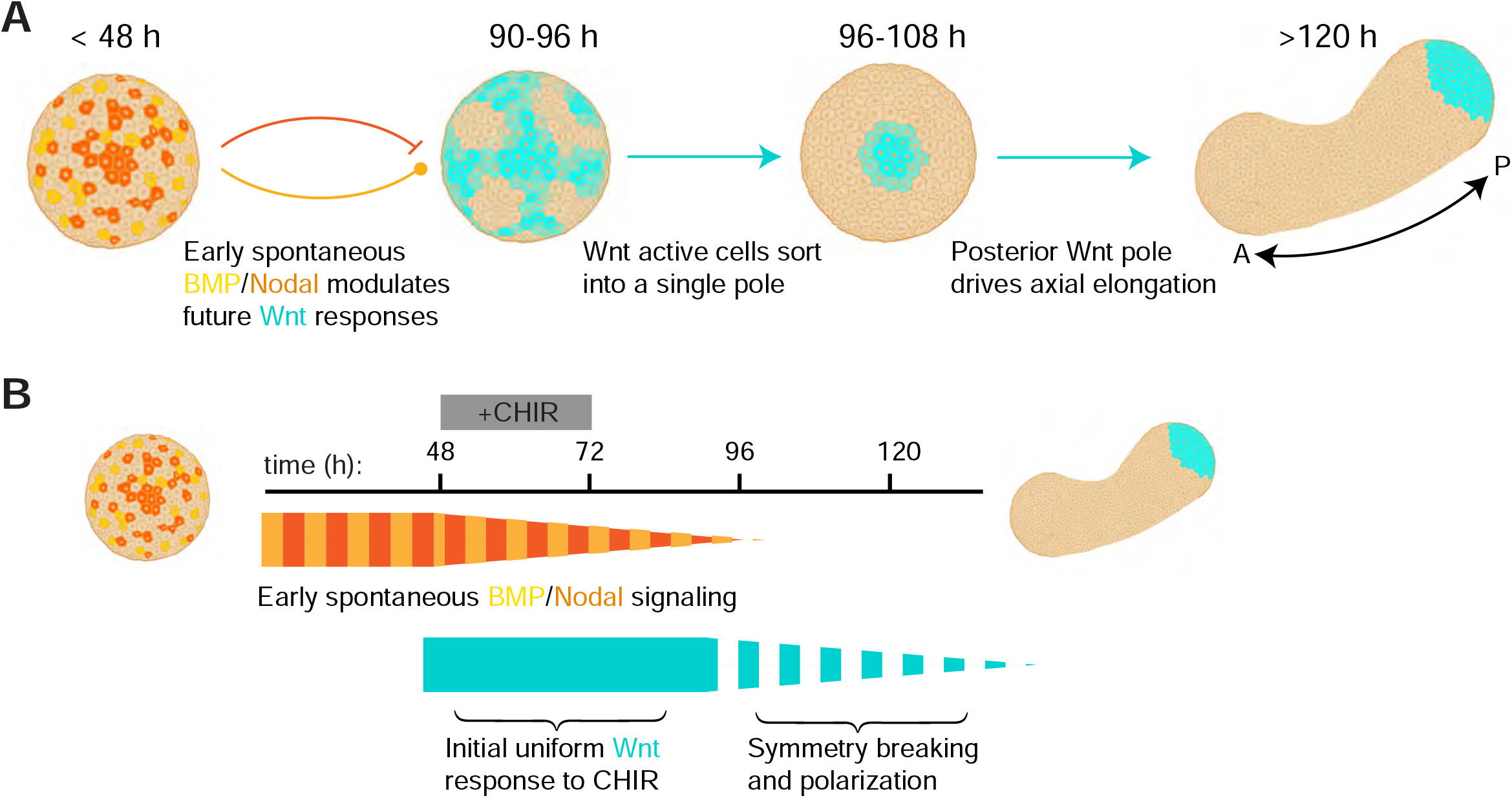
Proposed model of gastruloid symmetry breaking and polarization. **(A)** Illustration of the phases of symmetry breaking. Initial spontaneous Nodal/BMP activity modulates the response of cells to a uniform CHIR stimulus, yielding a heterogenous Wnt pattern. The heterogeneity ultimately resolves into a single pole of Wnt activity which organizes subsequent axial elongation. **(B)** Approximate timeline of signaling dynamics in model. Dashed bars indicate heterogeneity between cells; width of lines indicate relative fraction of signaling-active cells within the gastruloid.

How does this self-organizing model of axial morphogenesis relate to the program executed by the embryo? While the signaling pathways involved (Nodal, BMP and Wnt) appear to be largely conserved, their geometric organization is clearly different. In the mouse embryo, extra-embryonic sources of signaling ligands and inhibitors define separate anterior and posterior signaling compartments^39^. Studies in the zebrafish embryo have shown that spatially restricted sources of Nodal and BMP are sufficient to induce a complete secondary anterior-posterior axis^6^. Our results suggest that exogenous initial patterning of signaling domains is not strictly required to organize axial morphogenesis. The cell sorting dynamics we observe during gastruloid symmetry-breaking may represent a back-up or supplementary program which can refine cellular positions downstream of signaling gradients^8, 42^. Alternatively, it may play the role of an amplifier *in vivo* by converting weak initial asymmetries into persistent, all-or-none responses at the level of Wnt signaling activity, transcriptional identity, and cell position. Future studies examining the correspondence between gastruloid development and specific cell and tissue transformations in the embryo will be essential for further establishing the correct biological interpretation of gastruloid self-organization.

Our model also includes a potentially surprising role for Nodal signaling compared to prior studies in two-dimensional stem cell models. Whereas previous models suggest that it could either be a facilitator of Wnt signaling^40^, or a downstream effector with a BMP ◊ Wnt ◊ Nodal signaling hierarchy^43^, we find that spontaneous Nodal signaling can act upstream of Wnt to decrease the duration of stimulus response. While this regulatory role is not anticipated by the aforementioned models of Wnt and Nodal signaling, it is consistent with the Nodal’s role as a morphogen in the mouse embryo. By limiting Wnt activity, Nodal signaling patterns the anterior compartment of the gastruloid, just as it labels the anterior-most compartment of the primitive streak (i.e., the node)^39, 44^.

We emphasize that many components of our proposed model still await detailed investigation. For example, it is unknown how the initial heterogeneity in Nodal and BMP signaling activity is established; both stochastic gene expression and mechanical boundary conditions may play a role in setting up this heterogeneity. Many molecular connections also remain to be elucidated, such as the mechanistic relationship between prior TGFβ signaling and the duration of Wnt signaling after CHIR removal; to what extent specific adhesion molecules contribute to cell sorting; and to what extent cell movements are coordinated by long-range intracellular signals (e.g. Wnt inhibitors)^21^. Also, while Wnt signaling at 96 h is predictive of future cell positions and fates within the gastruloid, it is also possible that information is also specified by cells’ transcriptional states independently of Wnt. The relative contribution of Wnt-dependent and Wnt-independent information remains to be investigated.

Finally, we note that our study describes symmetry breaking in just one context – gastruloids prepared using culture conditions that initially exhibit homogeneous Wnt activity, and which recapitulate a posterior-biased (node-to-tail) portion of the mouse embryo’s A-P axis. Other routes to symmetry breaking may be observed in other contexts. For example, a recent study reported differences in Wnt activity between inner and outer cells in the gastruloid even during the CHIR pulse^16^, suggesting that Wnt symmetry-breaking has already occurred under those conditions. It may also be the case that the molecular logic of axial self-organization differs in more anterior compartments^45^ or in other model organisms^46, 47^. Addressing the processes at work in different cellular contexts will be essential to form a complete description of the molecular and physical processes underlying vertebrate axial polarization.

Through this work, we have deployed synthetic gene circuits to define the consequential signaling events for cells’ subsequent fates and positions. We demonstrated that three such circuits – Wnt-, Nodal-, and BMP-Recorder – are orthogonal to one another, capture information with time resolution < 6 h, and permit long-term tracing of subsequent cell populations by imaging and single-cell RNA sequencing. However, these signal-recording circuits are by no means the most sophisticated designs one may envision. A broad range of DNA-based signal recorders have been developed that encode signaling information as CRISPR-induced mutations or recombinase-encoded outcomes^48–50^. While the recombinase circuits implemented here sacrifice information bandwidth (each cell only implements two fluorescent states, for a total of one 1 bit of information), they excel in recording efficiency (nearly 100% of cells labeled during a 3 h recording) and temporal resolution (labeling in as little as 1 h). In the context of the gastruloid, additional information could also be gained by seeding mosaic gastruloids from multiple mESC lines that each contain a different recording circuit^51^. Pairs of split recombinases could also be used to realize complex logic or multiplexed recordings within single cells^52^. Eventually, optimized CRISPR-based recorders may enable high-bandwidth recording of a great many morphogen signals in parallel. We anticipate that molecular signal recording will represent a powerful strategy to decipher principles of biological self-organization.

## Methods

### Plasmids and cloning

Linear DNA fragments were amplified via PCR using CloneAmp HiFi PCR premix (Takara Bio, 639298). PCR products were cut from agarose gels and purified using the Nucleospin gel purification kit (Takara Bio, 240609). Plasmids were assembled from linear fragments using In-Fusion HD (Takara Bio, 638910) and amplified in Stellar chemically competent E. coli (Takara Bio, 636763) via ampicillin-resistant selection. Plasmid DNA was extracted by miniprep (Qiagen, 27104). All plasmid verification was performed by Sanger sequencing (Azenta) or nanopore sequencing (Plasmidsaurus).

Expression vectors were cloned into a piggyBac vector plasmid containing 5’ and 3’ flanking repeats to facilitate PBase insertion^28^. Plasmids encoding Cre recombinase, the ‘stoplight’ recording locus, and the Wnt-sensitive TOPFlash enhancer (P_TCF/LEF_) were purchased from Addgene (#89573, # 62732, and #12456, respectively). Sentinel enhancers responding to transcriptional output from Activin/Nodal (P_AR8_) and BMP (P_IBRE4_) were subcloned from AR8-Cerulean and IBRE4-Cerulean constructs generously shared by Kenneth Zaret. Plasmids generally contained a constitutively expressed fluorescent protein to serve as a selection marker during cell line generation.

### Cell culture

All reported experiments were performed using cell lines derived from E14tg2a mouse embryonic stem cells (ATCC CRL-1821). E14tg2a was first thawed and plated on mitotically inactivated feeder cells (Millipore Sigma PMEF-DR4-M) in basal growth media comprising GMEM (Millipore Sigma, G6148) supplemented with 10% ESC qualified fetal bovine serum (R&D Systems, S10250), 1× GlutaMAX (Gibco, 35050-061), 1× MEM non-essential amino acids (Gibco, 11140-470 050), 1 mM sodium pyruvate (Gibco, 11360-070), 100 μM 2-mercaptoethanol (Gibco, 471 21985-023), and 100 units/mL penicillin/streptomycin (Gibco, 15140-122). After reconstitution, E14tg2a cells were trypsinized and passaged onto a 25 cm^2^ tissue culture flask coated with 0.1% gelatin and grown ‘2i + LIF’ media comprising basal growth media further supplemented with 1000 units/mL LIF (Millipore 473 Sigma, ESG1107), 2 μM PD0325901 (Tocris, 4192), and 3 μM CHIR99021 (Tocris, 4423).

Propagation of cell lines was thereafter performed in 2i + LIF media on gelatin-coated tissue culture plastic unless otherwise noted. Cells were maintained between 20% and 75% confluency. Passaging was performed by aspirating remaining growth media, washing in phosphate buffered saline (Gibco, 14190144), and trypsinizing for 5 minutes at 37 C (TrypLE Express, Gibco, 12605028). Trypsin was quenched with 2i + LIF media, and cells were then pelleted by centrifugation at 135 rcf for 5 minutes. Residual trypsin and media were then aspirated and replaced with fresh media prior to replating in a fresh tissue culture vessel. Passage ratios varied from 1:5 to 1:10.

### Cell line generation

Engineered mESC lines were generated using the piggyBac random integration system. A chassis cell line was first grown to low-to-mid confluency (30-50%) in a 35 mm dish in 2i + LIF media. Transfection mixtures were prepared in 250 µL Gibco OptiMEM (Fisher Scientific 31-985-070) and further comprised 5 µL Lipofectamine STEM transfection reagent (ThermoFisher STEM00001), 2080 ng vector plasmid, and 420 ng PBase ‘helper’ plasmid (System Biosciences). After preparation, transfection mixtures were equilibrated at room temperature for 30 minutes. Immediately preceding transfection, chassis cell cultures were then given fresh 2i + LIF media. Transfection mixture was then added dropwise to chassis cells, and gently mixed via rocking.

Cultures were propagated for at least 4 days following transfection, by which vector expression is predominantly driven by genomically-integrated constructs. Cells were then trypsinized into a single-cell suspension for fluorescently activated cell sorting (Sony SH800). Cytometry events were first sorted for single cells based on scattering profiles, and then transformed cells were identified via fluorescent signals. Candidate clonal cultures were generated by sorting single cells into separate wells of a 96-well plate. Following sorting, colonies remained unperturbed for 7-10 days, after which they were assessed via microscopy to identify promising candidates via fluorescent intensity and colony morphology (i.e. preferring round colonies with smooth boundaries). Candidate clonal colonies were then identified and supplemented with an additional 150 µL of 2i + LIF media, and fed every other day until 14 days post-seeding. At 14 days, promising clones were trypsinized and passaged onto a 12-well plate. Lines which remained viable were then further expanded and functionally assessed.

### Signal recording benchmarking

Signal recording cell lines were benchmarked in adherent cultures grown on gelatin-coated tissue culture plastic. Minimum recording window measurements (**Figure 2C**) and doxycycline concentration calibration (**Figure S2A**) were performed in steady-state 2i + LIF media. All other measurements were performed in N2B27 basal media, supplemented with doxycycline, CHIR, or recombinant morphogen proteins according to the experimental condition. Media was exchanged to fresh N2B27 at least 24 h prior to recording experiments to allow cells to equilibrate to basal media conditions. For Wnt recorder fidelity testing (**Figure 2C**), Wnt dose response (**Figure S2B**), and morphogen crosstalk experiments (**Figure 2E**), morphogen signals were added to media simultaneously with the onset of the doxycycline recording window, and then washed out following cessation of the doxycycline recording window. Cells were expanded for at least 24 h following cessation of doxycycline treatment and then assayed via flow cytometry.

### Gastruloid protocol

Gastruloids were grown in N2B27 media comprising 1:1 mixture of DMEM/F-12 (Gibco, 11320033) and neurobasal medium (Gibco, 21103049), supplemented with 100 μM 2-mercaptoethanol, 1:100 N-2 (Gibco, 17502048), 1:50 B-27 (Gibco, 17504044), and 100 units/mL penicillin/streptomycin (Gibco, 15140-122). Seed cultures were maintained in 2i + LIF media prior to gastruloid formation to minimize pre-existing heterogeneity in early stage gatruloids which may bias symmetry breaking. For ‘LIF only’ preculture gastruloids, cells were transferred from 2i LIF media to media without either PD0325901 or CHIR99021 for 6 days (2 passages) prior to gastruloid formation.

To form gastruloids, seed cultures were trypsinized, pelleted, and then washed twice with phosphate-buffered saline (separated by additional 5 minute centrifugations at 135 rcf) to remove residual CHIR 99021. Following the second PBS wash, cells were resuspended in N2B27 media and transferred to the cell sorter for fluorescence-activated cell sorting. Sorting events were gated for single cells based on scattering profiles. For signal recording gastruloids, cells were further gated against GFP expression to avoid contamination from the small fraction of cells (∼0.1%) which had already excised DsRed. To form gastruloids, 200 single cells were sorted into each of the 60 central wells of a 96-well round-bottom ultra-low attachment microplates (Corning, 7007). Central wells were pre-filled with 40 µL of N2B27 to receive cells. To minimize the effects of evaporation at the edges of the plate, the perimeter wells were filled with 150 μL PBS supplemented with 1000 units/mL penicillin/streptomycin.

Following seeding, gastruloids were transferred to a cell culture incubator and left unperturbed for 48 h unless otherwise noted. At t = 48 h, gastruloids were fed with 150 uL of N2B27 media further supplemented with 3 uM CHIR to stimulate Wnt activity. At t = 72 hours, 150 µL of CHIR-containing media was removed from gastruloid wells, and then the gastruloids were fed with 150 µL of fresh N2B27 media without CHIR. A similar feeding (150 µL media remoed, 150 µL fresh N2B27 added) was performed every 24 hours for the remained of gastruloid morphogenesis. Signal recording during gastruloid morphogenesis was performed via the transient addition of 150 µL media supplemented with 100 ng/mL doxycycline. Recording windows were either 90 min (for Wnt recording) or 3 h (for BMP and Nodal). Doxycycline was washed out with two successively 150 µL media changes, leading to a total dilution of approximately 1:20 (sufficient to bring the doxycline concentraton below the recording threshold defined in **Figure S2A**).

Gastruloids were assayed either by imaging or by flow cytometry. To prepare samples for imaging, gastruloids were fixed for 2 h in 4% PFA at 4 °C on a nutator. Samples were then washed twice in PBS to remove PFA, and either transferred for immunofluorescence labeling or immediately transferred to a glass-bottom 96 well plate for imaging. To assess gastruloids via flow cytometry, gastruloids were first pooled into a 1.5 mL Eppendorf tube, centrifugated (500 rcf, 3 min), washed in PBS, and then trypsinized in 100 µL of TrypLE Express at 37 °C. Gastruloids were trypsinized for 3 minutes, retrieved and triturated to dislodge gastruloids, and returned to 37 for an additional 3 minutes to complete digestion. Trypsinization was then quenched with 300 µL of N2B27, after which samples were immediately assayed via flow cytometry.

### Immunofluorescence

Gastruloid staining was performed according to previously reported protocols^17^. Briefly, cohorts of PFA fixed gastruloids were pooled into single wells of ultra low-attachment 96 well plates and washed twice with PBS to remove any residual PBS. All buffer exchanges were performed under a dissection microscope to maximize buffer turnover while minimizing sample loss due to accidental aspiration. Gastruloids were permeabilized overnight at 4 C with nutation in PBSFT buffer (89.8% phosphate buffered saline, 10% fetal bovine serum, and 0.2% Triton X-100). PBSFT was then removed and exchanged for PBSFT containing dilutions of primary antibodies against targets of interest. Primary dilutions used were: 1:200 for rabbit anti-Pcdh19 (abcam ab191198); 1:100 for rabbit anti-Aldh1a2 (abcam ab156019); and 1:200 for rabbit anti-Cdh1 (Cell Signaling Technology 3195T). Primary antibody incubation was performed overnight at 4 C with nutation. Following primary incubations, samples were treated with PBSFT for three consecutive washes. Additional 3× serial wash sequences were performed two additional times, with 1 hour of nutation spacing each wash sequence, for 9 total washes. Samples were then incubated with fluorescently conjugated secondary antibodies (Goat anti-rabbit Alexa Fluor 647 conjugate, Invitrogen A27040) diluted 1:500 in PBSFT overnight at 4 °C with nutation. Following secondary antibody incubation, samples were again washed nine times following the same protocol described above. Samples were then transferred to a glass bottom 96-well plate for imaging (Cellvis P96-1.5H-N).

### Image processing and quantification

All imaging data reported were acquired on a Nikon Eclipse Ti confocal microscope with a 600 Prior linear motorized stage, a Yokogawa CSU-X1 spinning disk, an Agilent laser line module 601 containing 405, 488, 561 and 650nm lasers, and an iXon DU897 EMCCD camera. Images were acquired as three-dimensional hyperstacks using Nikon elements and converted to maximum intensity projections in ImageJ. For low-signal regimes (P_TCF/LEF_-GFP live imaging, late stage gastruloid P_TCF/LEF_-iRFP patterns) median filtering was performed (2 to 5 pixel kernel) to denoise and improve contrast. Near-infrared images were further background subtracted to correct for a heterogeneous background field using a sample-free background image.

Quantification was performed using custom software in MATLAB. Briefly, maximum intensity projections were normalized on a [0,1] interval by first subtracting background fluorescence and then dividing by the 99^th^ percentile pixel value, each on a channel-by-channel basis. For sparse images (e.g. sparse GFP labeling), the maximum pixel percentile was increased to avoid over-amplification. Gastruloid segmentation was performed on a morphological image generated by adding 488 nm and 561 nm excitation channel images (to account for total recorder locus signal). Morphological images were binarized; manually trimmed to separate neighboring gastruloids from co-segmenting; and then dilated, eroded, and filled to smoothen boundaries.

Individual gastruloids were linearized by first skeletonizing with the MATLAB bwskel() function (MinBranchLength = 200). The resultant anterior-posterior axis trace was clipped if necessary (to avoid over-fitting on boundary artifacts), and manually extended to the anterior and posterior poles to create a curvilinear axis. The posterior pole of the axis was manually identified based on gastruloid morphology and/or Wnt signaling activity, and the curvilinear axis was assigned units of distance based on geodesic distance from the anterior pole computed by bwdistgeodesic(). Pixels contained within a gastruloid segment were assigned an axial coordinate based on the geodesic distance of the closest curvilinear axis point, and then binned over intervals of dL = 10 microns. One-dimensional fluorescence profiles were computed by averaging values within all gastruloid pixels corresponding to a single axial distance bin.

For Wnt recording quantification, fractional labeling profiles are reported as the ratio of the average normalized GFP fluorescence to the sum of the average normalized GFP and average normalized DsRed fluorescence. To average profiles over experimental replicates, profiles were first rescaled onto a unit anterior-posterior axis x/L = [0,1] with bins corresponding to 1% of the overall axis length using linear interpolation. For each experimental condition, both mean values and standard deviations were computed for each axial position. To compute the integrated Wnt signaling activity:

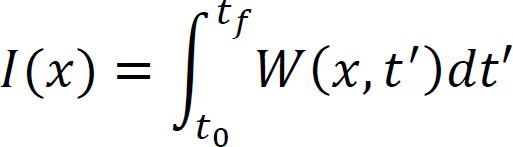

(**Figure 3F**), fractional labeling profiles were first individually normalized to unit integral over the A-P axis and then added from t_dox_ = 96 h to t_dox_ = 120 h. An additional profile from t_f_ = 134 h was extracted from final P_TCF/LEF_-iRFP fluorescence and similarly normalized to unit integral along the A-P axis, and then added to the total integral *I(x)* to capture the final signaling profile.

Spatial profiles of Wnt signaling in early gastruloids (**Figure S1F**) were quantified to assess relative heterogeneity and polarization (**Figure 1G**). Gastruloids were first normalized and segmented as described above. Images were then further denoised using a median filter (14 um × 14 um kernel) to remove high spatial frequency noise. Relative heterogeneity was determined by computing the standard deviation of pixel intensities within a segmented gastruloid area.

Relative polarization was computed according to

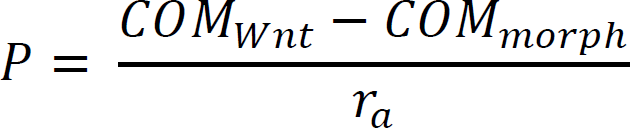

Where *COM_morph_* is the morphological center of mass (determined by EF1a-DsRed fluorescence) and *COM_MWnt_* is the center of mass of the Wnt activity pattern, and *r_a_* is the morphological semimajor axis length.

### Live Imaging

Gastruloids were immobilized for live imaging via Matrigel embedding. To prepare Matrigel matrix (Corning #356231), frozen aliquots were thawed overnight at 4 °C. Glass-bottom 35 mm dishes (Cellvis D35-20-1.5-N) and pipette tips were chilled to temperature controlled cold room at 4 C, and the central imaging well was then coated with Matrigel reagent. The Matrigel-coated dish was then transferred to a metal surface on ice to prevent gelling. Gastruloids were then collected and dispersed into cooled Matrigel while maintaining minimal media carryover (roughly 10 µL volume). Approximately 6 to 12 gastruloids were embedded per experiment. Gastruloids were manually separated via pipette manipulation to ensure even spacing, and then transferred to 37 °C for 10 minutes to solidify the Matrigel matrix. 2 mL of prewarmed N2B27 medium was then added to the glass-bottom dish. The sample was then transferred to a microscope-mounted environmental chamber (Okolab) to maintain appropriate temperature, humidity, and CO_2_ during imaging. Samples were imaged using a Nikon Eclipse Ti confocal microscope. Movies were collected over 24 to 30 h time periods, with frames acquired once every 10 minutes. Individual frames were acquired at 4 z-planes. Images were median filtered to denoise and maximum-intensity projected to generate final movies.

### Single cell RNA sequencing

To prepare gastruloids for single cell sequencing, samples were first pooled into a 1.5 mL Eppendorf tube. At least 50 gastruloids were pooled for each experimental condition. Following pooling, gastruloids were centrifugated (500 rcf for 3 minutes), washed in PBS, re-centrifugated (500 rcf for 3 minutes), and then resuspended in 100 µL trypsin and transferred to 37°C. After 3 minutes, the gastruloid-trypsin mixture was retrieved, triturated to dislodge gastruloids, and then returned to 37°C for an additional 3 minutes. Samples were then quenched with 300 µL N2B27 media, and immediately transferred to a cell sorter (Sony SH800). For 96-hour gastruloids, cytometer events corresponding to single cells were identified based on light scattering profiles and then recovered into 500 µL of N2B27 media. For 120-hour gastruloids, single cell events were further gated based on GFP fluorescence and recovered into two separate recovery tubes (each containing 500 µL N2B27 media). Recovered cells were immediately centrifugated (500 rcf for 5 min), and then resuspended in 50 µL of PBS. A 10 µL fraction was then stained for dead cells with trypan blue and automatically counted (Invitrogen Countess) to assess density and viability of recovered cells. Samples were then diluted to a target cell density of 1000 cells/µL. Both 96-hour and 120-hour samples maintained over 80% cell viability by this step.

Following dilution in PBS, cells were kept on ice and transferred to a Chromium Controller (10× Genomics) to generate gel-in-bead emulsions (GEM) for single cell RNA labeling. Microfluidic chip lanes (Chip K, 10× Genomics) were loaded to a target recovery of 5000 cells/lane. We used the Single Cell 5’ v2 Reagent Kit (10× Genomics) to enable optional direct capture of non-polyadenylated mRNAs from synthetic reporter genes. Following GEM formation, the RT reaction was performed with mastermix supplemented with direct capture primer targeting iRFP (5 pmol per sample lane):

5’-AAGCAGTGGTATCAACGCAGAGTACCTCTTCCATCACGCCGATCTG – 3’

Gene expression (GEx) libraries were then generated from sample cDNA according to manufacturer protocol. Briefly, cDNA samples were fragmented and selected for a target size via double-sized magnetic bead selection (Beckman-Coulter SPRIselect). Samples were then ligated with adaptor oligos and PCR-amplified with dual library indices for sample demultiplexing. Library concentrations were assessed by fluorometry (Qubit) and size distributions were measured with automated electrophoresis (Agilent Bioanalyzer). Samples were pooled to target equal concentration and submitted for pooled sequencing on an Illumina NovaSeq SP 100nt Flowcell v1.5. The library was sequenced to a depth of 47,000 reads/cell.

### Single cell RNA sequencing data analysis

Raw FastQ files were demultiplexed by Dr. Wei Wang at the Princeton Genomics Core Facility. The Mm10_2020 transcriptome was modified to include BFP, iCre, iRFP, rTTA, and the “Stoplight Recorder” as additional elements for alignment in the analysis pipeline, as per 10× documentation. Demultiplexed files were then individually converted to counts tables using the 10× CellRanger 6.0.1 Pipeline per outlined documentation. All jobs were submitted through the Princeton LSI gencomp2 computing cluster.

Single cell mRNA count matrices were analyzed with the Single cell analysis in Python (Scanpy) toolkit. For 120 h datasets, GFP-positive and GFP-negative libraries were conjoined into a single dataframe retaining conditional labels. Initial filtering was performed to exclude genes detected in fewer than 3 cells, and cells with either fewer than 500 genes or fewer than 3000 total assigned reads. Secondary filtering was performed to exclude cells with either greater than 8% counts corresponding to mitochondrial genes or greater than 30% counts corresponding to ribosomal RNA. Counts were then normalized to 10,000 counts/cell, log-transformed, regressed to remove variation contributed by cell cycle-associated genes. Highly variable genes were then normalized to zero mean and unit variance, batch corrected, and decomposed via PCA. A nearest-neighbor graph was computed on the top 40 principal components using a 20-member local neighborhood, and visualized by Uniform Manifold Approximation and Projection (UMAP). Cell types were identified by Leiden clustering (resolution = 0.4 for 96 h data; resolution = 0.6 for 120 h data).

Leiden clusters were associated with previously annotated reference cell types by comparing differentially expressed marker genes between both datasets. Mean logarithmized expression of genes within each cluster was compared to overall mean via t-test. Candidate marker genes for each cluster were ranked according to test statistics and filtered for p-values < 0.01, and then up to 50 top ranked genes were retained as marker genes for each Leiden cluster. Marker genes were then compared to a reference atlas^32^ comprising 30 marker genes for previously annotated embryonic cell types to determine overlap of shared markers between Leiden clusters and reference atlas cell types. The statistical likelihoods of observed overlaps were computed by comparing overlap degrees to the binomial probability of drawing the same number of common genes from a random list of all unique atlas marker genes.

Signal recording fractions within Leiden clusters were calculated by comparing relative distributions of cell types across recording conditions. For each cluster *k*, we calculated the conditional probability that a cell would occupy cluster *k* given its recording condition *g*_±_ as:

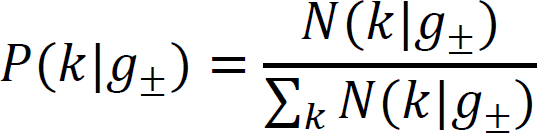

An overall relative labeling statistic for each cluster was then computed by comparing conditional probabilities across recording values:

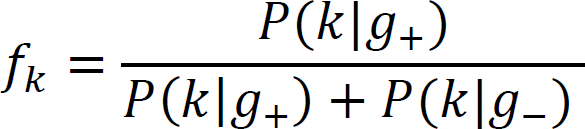

## Author Contributions

Conceptualization, H.M.M., M.M.C., and J.E.T.; Methodology, H.M.M., S.S., M.M.C., and J.E.T.; Investigation, H.M.M.; Funding, H.M.M., J.E.T.; Writing and Editing, H.M.M. and J.E.T.; Supervision, J.E.T., B.A.

## Acknowledgements

The authors thank all members of the Toettcher lab, particularly Evan Underhill, for insightful comments and suggestions. We also thank Alfonso Martinez-Arias for helpful discussions and feedback on the manuscript; Ken Zaret for sharing reporter plasmid constructs; Aaron Lin for advice on single-cell sequencing; and the Lewis Sigler Institute Genomics Core Facility team for technical support (Wei Wang, Jennifer Miller, and Jean Arly Volmar). This work was supported by Lewis Sigler Scholars program and the NSF Center for the Physics of Biological Function PHY1734030 (H.M.M.); NIH grant T32GM007388 (S.C.S.); NSF RECODE grant 2134935, NIH grant U01DK127429, and a Vallee Scholars award (J.E.T.).

## Competing Interests

J.E.T. is a scientific advisor for Prolific Machines and Nereid Therapeutics. B.A. is an advisory board member for Arbor Biotechnologies and Tessera Therapeutics and holds equity in Celsius Therapeutics. H.M.M. is a cofounder and scientific advisor for C16 Biosciences. The remaining authors declare no conflicts of interest.

## References

1. Spemann, H., and Mangold, H. (1924). über Induktion von Embryonalanlagen durch Implantation artfremder Organisatoren. Archiv f mikr Anat u Entwicklungsmechanik 100, 599– 638. 10.1007/BF02108133.

2. Fauny, J.-D., Thisse, B., and Thisse, C. (2009). The entire zebrafish blastula-gastrula margin acts as an organizer dependent on the ratio of Nodal to BMP activity. Development 136, 3811–3819. 10.1242/dev.039693.

3. Turing, A.M. (1952). The Chemical Basis of Morphogenesis. Philosophical Transactions of the Royal Society of London. Series B, Biological Sciences 237, 37–72.

4. Kondo, S., and Miura, T. (2010). Reaction-Diffusion Model as a Framework for Understanding Biological Pattern Formation. Science 329, 1616–1620. 10/bhd5cf.

5. Muller, P., Rogers, K.W., Jordan, B.M., Lee, J.S., Robson, D., Ramanathan, S., and Schier, A.F. (2012). Differential Diffusivity of Nodal and Lefty Underlies a Reaction-Diffusion Patterning System. Science 336, 721–724. 10.1126/science.1221920.

6. Xu, P.-F., Houssin, N., Ferri-Lagneau, K.F., Thisse, B., and Thisse, C. (2014). Construction of a Vertebrate Embryo from Two Opposing Morphogen Gradients. Science 344, 87–89. 10.1126/science.1248252.

7. Toda, S., Blauch, L.R., Tang, S.K.Y., Morsut, L., and Lim, W.A. (2018). Programming self-organizing multicellular structures with synthetic cell-cell signaling. Science 361, 156–162. 10.1126/science.aat0271.

8. Tsai, T.Y.-C., Sikora, M., Xia, P., Colak-Champollion, T., Knaut, H., Heisenberg, C.-P., and Megason, S.G. (2020). An adhesion code ensures robust pattern formation during tissue morphogenesis. Science 370, 113–116. 10.1126/science.aba6637.

9. Hannezo, E., and Heisenberg, C.-P. (2019). Mechanochemical Feedback Loops in Development and Disease. Cell 178, 12–25. 10.1016/j.cell.2019.05.052.

10. Turner, D.A., Girgin, M., Alonso-Crisostomo, L., Trivedi, V., Baillie-Johnson, P., Glodowski, C.R., Hayward, P.C., Collignon, J., Gustavsen, C., and Serup, P. (2017). Anteroposterior polarity and elongation in the absence of extra-embryonic tissues and of spatially localised signalling in gastruloids: mammalian embryonic organoids. Development 144, 3894–3906.

11. Beccari, L., Moris, N., Girgin, M., Turner, D.A., Baillie-Johnson, P., Cossy, A.-C., Lutolf, M.P., Duboule, D., and Arias, A.M. (2018). Multi-axial self-organization properties of mouse embryonic stem cells into gastruloids. Nature 562, 272–276. 10.1038/s41586-018-0578-0.

12. van den Brink, S.C. Alemany, A., van Batenburg, V., Moris, N., Blotenburg, M., Vivié, J., Baillie-Johnson, P., Nichols, J., Sonnen, K.F., Martinez Arias, A., et al. (2020). Single-cell and spatial transcriptomics reveal somitogenesis in gastruloids. Nature 582, 405–409. 10.1038/s41586-020-2024-3.

13. Sheng, G., Martinez Arias, A., and Sutherland, A. (2021). The primitive streak and cellular principles of building an amniote body through gastrulation. Science 374, abg1727. 10.1126/science.abg1727.

14. Beddington, R.S. (1994). Induction of a second neural axis by the mouse node. Development 120, 613–620. 10.1242/dev.120.3.613.

15. Baillie-Johnson, P., Brink, S.C. van den Balayo, T., Turner, D.A., and Arias, A.M. (2015). Generation of Aggregates of Mouse Embryonic Stem Cells that Show Symmetry Breaking, Polarization and Emergent Collective Behaviour In Vitro. JoVE (Journal of Visualized Experiments), e53252. 10.3791/53252.

16. Suppinger, S., Zinner, M., Aizarani, N., Lukonin, I., Ortiz, R., Azzi, C., Stadler, M.B., Vianello, S., Palla, G., Kohler, H., et al. (2023). Multimodal characterization of murine gastruloid development. Cell Stem Cell. 10.1016/j.stem.2023.04.018.

17. Underhill, E.J., and Toettcher, J.E. (2023). Control of gastruloid patterning and morphogenesis by the Erk and Akt signaling pathways. bioRxiv.

18. Hashmi, A., Tlili, S., Perrin, P., Lowndes, M., Peradziryi, H., Brickman, J.M., Arias, A.M., and Lenne, P.-F. (2022). Cell-state transitions and collective cell movement generate an endoderm-like region in gastruloids. Elife 11, e59371.

19. Petersen, C.P., and Reddien, P.W. (2009). Wnt Signaling and the Polarity of the Primary Body Axis. Cell 139, 1056–1068. 10.1016/j.cell.2009.11.035.

20. Anlaş, K., Gritti, N., Oriola, D., Arató, K., Nakaki, F., Lim, J.L., Sharpe, J., and Trivedi, V. (2021). Dynamics of anteroposterior axis establishment in a mammalian embryo-like system. 2021.02.24.432766. 10.1101/2021.02.24.432766.

21. Anand, G.M., Megale, H.C., Murphy, S.H., Weis, T., Lin, Z., He, Y., Wang, X., Liu, J., and Ramanathan, S. (2023). Controlling organoid symmetry breaking uncovers an excitable system underlying human axial elongation. Cell 186, 497–512.e23. 10.1016/j.cell.2022.12.043.

22. Liu, P., Wakamiya, M., Shea, M.J., Albrecht, U., Behringer, R.R., and Bradley, A. (1999). Requirement for Wnt3 in vertebrate axis formation. Nat Genet 22, 361–365. 10.1038/11932.

23. Korinek, V., Barker, N., Morin, P.J., van Wichen, D., de Weger, R., Kinzler, K.W., Vogelstein, B., and Clevers, H. (1997). Constitutive Transcriptional Activation by a β-Catenin-Tcf Complex in APC−/− Colon Carcinoma. Science 275, 1784–1787. 10.1126/science.275.5307.1784.

24. Ying, Q.L., Wray, J., Nichols, J., Batlle-Morera, L., Doble, B., Woodgett, J., Cohen, P., and Smith, A. (2008). The ground state of embryonic stem cell self-renewal. Nature 453, 519–523. 10.1038/nature06968.

25. Kretzschmar, K., and Watt, F.M. (2012). Lineage Tracing. Cell 148, 33–45. 10.1016/j.cell.2012.01.002.

26. van de Moosdijk, A.A.A., van de Grift, Y.B.C., de Man, S.M.A., Zeeman, A.L., and van Amerongen, R. (2020). A novel Axin2 knock-in mouse model for visualization and lineage tracing of WNT/CTNNB1 responsive cells. Genesis 58, e23387. 10.1002/dvg.23387.

27. Reijmers, L.G., Perkins, B.L., Matsuo, N., and Mayford, M. (2007). Localization of a Stable Neural Correlate of Associative Memory. Science 317, 1230–1233. 10.1126/science.1143839.

28. Yusa, K., Zhou, L., Li, M.A., Bradley, A., and Craig, N.L. (2011). A hyperactive piggyBac transposase for mammalian applications. Proc Natl Acad Sci U S A 108, 1531–1536. 10.1073/pnas.1008322108.

29. Serup, P., Gustavsen, C., Klein, T., Potter, L.A., Lin, R., Mullapudi, N., Wandzioch, E., Hines, A., Davis, A., Bruun, C., et al. (2012). Partial promoter substitutions generating transcriptional sentinels of diverse signaling pathways in embryonic stem cells and mice. Disease Models & Mechanisms 5, 956–966. 10.1242/dmm.009696.

30. Das, A.T., Tenenbaum, L., and Berkhout, B. (2016). Tet-On Systems For Doxycycline-inducible Gene Expression. Curr Gene Ther 16, 156–167. 10.2174/1566523216666160524144041.

31. Yang, Y.S., and Hughes, T.E. (2001). Cre stoplight: a red/green fluorescent reporter of Cre recombinase expression in living cells. Biotechniques 31, 1036, 1038, 1040–1041. 10.2144/01315st03.

32. Grosswendt, S., Kretzmer, H., Smith, Z.D., Kumar, A.S., Hetzel, S., Wittler, L., Klages, S., Timmermann, B., Mukherji, S., and Meissner, A. (2020). Epigenetic regulator function through mouse gastrulation. Nature 584, 102–108. 10.1038/s41586-020-2552-x.

33. van den Brink, S.C., Alemany, A., van Batenburg, V., Moris, N., Blotenburg, M., Vivié, J., Baillie-Johnson, P., Nichols, J., Sonnen, K.F., Martinez Arias, A., et al. (2020). Single-cell and spatial transcriptomics reveal somitogenesis in gastruloids. Nature 582, 405–409. 10.1038/s41586-020-2024-3.

34. Niederreither, K., Subbarayan, V., Dollé, P., and Chambon, P. (1999). Embryonic retinoic acid synthesis is essential for early mouse post-implantation development. Nat Genet 21, 444–448. 10.1038/7788.

35. Yaman, Y.I., and Ramanathan, S. (2023). Controlling human organoid symmetry breaking reveals signaling gradients drive segmentation clock waves. Cell 186, 513–527.e19. 10.1016/j.cell.2022.12.042.

36. Chal, J., and Pourquié, O. (2017). Making muscle: skeletal myogenesis in vivo and in vitro. Development 144, 2104–2122. 10.1242/dev.151035.

37. Steinberg, M.S. (1962). On the mechanism of tissue reconstruction by dissociated cells, i. population kinetics, differential adhesiveness, and the absence of directed migration*. Proceedings of the National Academy of Sciences 48, 1577–1582. 10.1073/pnas.48.9.1577.

38. Tsai, T.Y.-C., Garner, R.M., and Megason, S.G. (2022). Adhesion-Based Self-Organization in Tissue Patterning. Annual Review of Cell and Developmental Biology 38, 349–374. 10.1146/annurev-cellbio-120420-100215.

39. Bardot, E.S., and Hadjantonakis, A.-K. (2020). Mouse gastrulation: coordination of tissue patterning, specification and diversification of cell fate. Mech Dev 163, 103617. 10.1016/j.mod.2020.103617.

40. Massey, J., Liu, Y., Alvarenga, O., Saez, T., Schmerer, M., and Warmflash, A. (2019). Synergy with TGFβ ligands switches WNT pathway dynamics from transient to sustained during human pluripotent cell differentiation. Proc Natl Acad Sci U S A 116, 4989–4998. 10.1073/pnas.1815363116.

41. Camacho-Aguilar, E., Yoon, S., Ortiz-Salazar, M.A., and Warmflash, A. (2022). Combinatorial interpretation of BMP and WNT allows BMP to act as a morphogen in time but not in concentration. 2022.11.11.516212. 10.1101/2022.11.11.516212.

42. Pinheiro, D., Kardos, R., Hannezo, É., and Heisenberg, C.-P. (2022). Morphogen gradient orchestrates pattern-preserving tissue morphogenesis via motility-driven unjamming. Nat. Phys. 18, 1482–1493. 10.1038/s41567-022-01787-6.

43. Chhabra, S., Liu, L., Goh, R., Kong, X., and Warmflash, A. (2019). Dissecting the dynamics of signaling events in the BMP, WNT, and NODAL cascade during self-organized fate patterning in human gastruloids. PLOS Biology 17, e3000498. 10.1371/journal.pbio.3000498.

44. Conlon, F.L., Lyons, K.M., Takaesu, N., Barth, K.S., Kispert, A., Herrmann, B., and Robertson, E.J. (1994). A primary requirement for nodal in the formation and maintenance of the primitive streak in the mouse. Development 120, 1919–1928. 10.1242/dev.120.7.1919.

45. Girgin, M.U., Broguiere, N., Mattolini, L., and Lutolf, M.P. (2021). Gastruloids generated without exogenous Wnt activation develop anterior neural tissues. Stem Cell Reports 16, 1143–1155. 10.1016/j.stemcr.2021.03.017.

46. Cheng, T., Xing, Y.-Y., Liu, C., Li, Y.-F., Huang, Y., Liu, X., Zhang, Y.-J., Zhao, G.-Q., Dong, Y., Fu, X.-X., et al. (2023). Nodal coordinates the anterior-posterior patterning of germ layers and induces head formation in zebrafish explants. Cell Rep 42, 112351. 10.1016/j.celrep.2023.112351.

47. Fulton, T., Trivedi, V., Attardi, A., Anlas, K., Dingare, C., Arias, A.M., and Steventon, B. (2020). Axis Specification in Zebrafish Is Robust to Cell Mixing and Reveals a Regulation of Pattern Formation by Morphogenesis. Curr Biol 30, 2984–2994.e3. 10.1016/j.cub.2020.05.048.

48. Choi, J., Chen, W., Minkina, A., Chardon, F.M., Suiter, C.C., Regalado, S.G., Domcke, S., Hamazaki, N., Lee, C., Martin, B., et al. (2022). A time-resolved, multi-symbol molecular recorder via sequential genome editing. Nature 608, 98–107. 10.1038/s41586-022-04922-8.

49. Chen, W., Choi, J., Nathans, J.F., Agarwal, V., Martin, B., Nichols, E., Leith, A., Lee, C., and Shendure, J. (2021). Multiplex genomic recording of enhancer and signal transduction activity in mammalian cells. bioRxiv, 2021–11.

50. Bhattarai-Kline, S., Lear, S.K., Fishman, C.B., Lopez, S.C., Lockshin, E.R., Schubert, M.G., Nivala, J., Church, G.M., and Shipman, S.L. (2022). Recording gene expression order in DNA by CRISPR addition of retron barcodes. Nature 608, 217–225. 10.1038/s41586-022-04994-6.

51. Wehmeyer, A.E., Schüle, K.M., Conrad, A., Schröder, C.M., Probst, S., and Arnold, S.J. (2022). Chimeric 3D gastruloids – a versatile tool for studies of mammalian peri-gastrulation development. Development 149, dev200812. 10.1242/dev.200812.

52. Weinberg, B.H., Pham, N.T.H., Caraballo, L.D., Lozanoski, T., Engel, A., Bhatia, S., and Wong, W.W. (2017). Large-scale design of robust genetic circuits with multiple inputs and outputs for mammalian cells. Nat Biotechnol 35, 453–462. 10.1038/nbt.3805.

