## Supplementary material for "Recording morphogen signals reveals origins of gastruloid symmetry breaking": SupplmentaryInfo

**This PDF file includes:**

Supplementary Figures 1-6
Supplementary Movie Legends

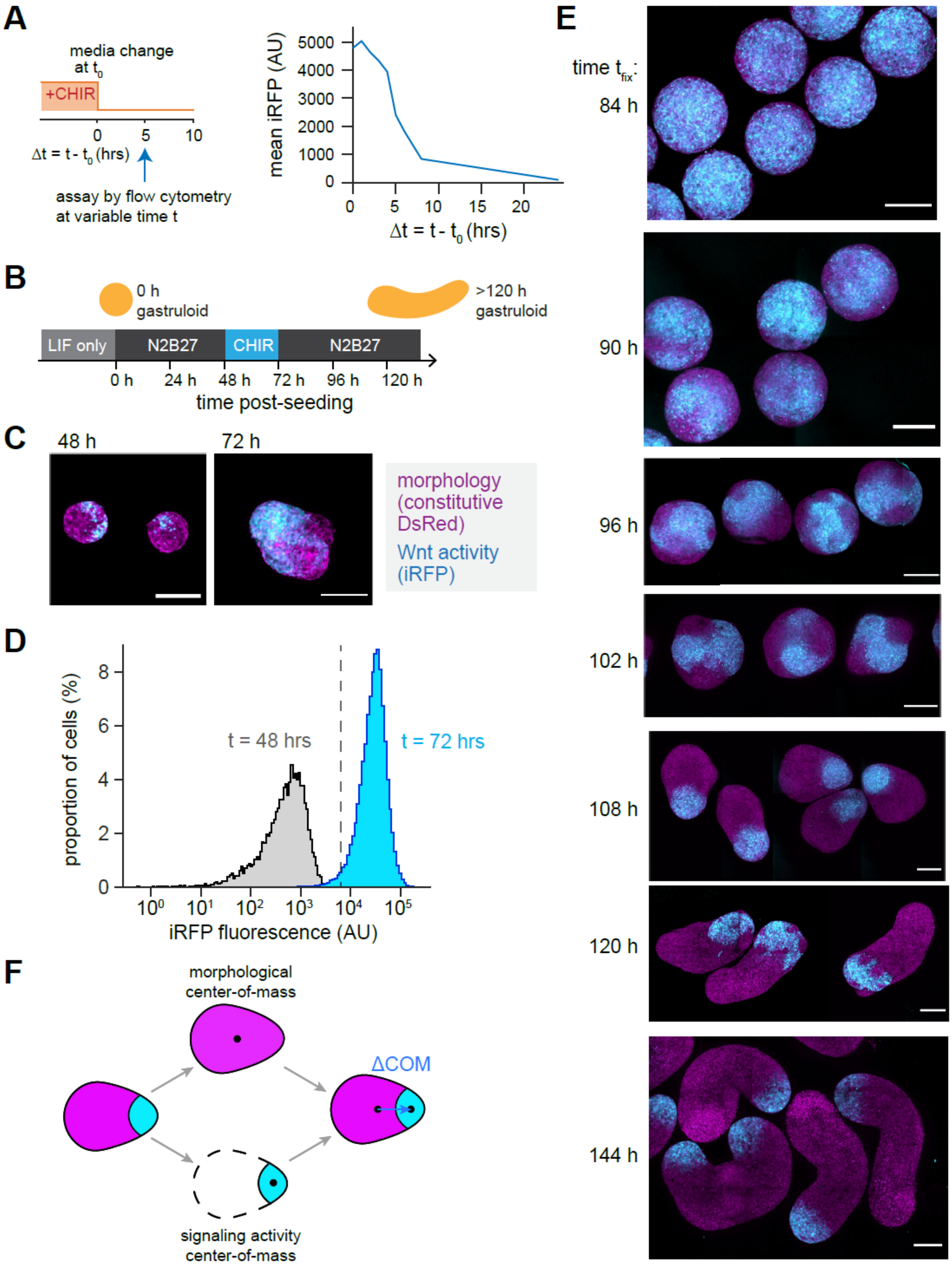

A
media change
at $t_0$
+CHIR
0
5
10
Δt = t - $t_0$ (hrs)
assay by flow cytometry
at variable time t
mean iRFP (AU)
5000
4000
3000
2000
1000
0
0
5
10
15
20
Δt = t - $t_0$ (hrs)
B
0 h
gastruloid
>120 h
gastruloid
LIF only
N2B27
CHIR
N2B27
0 h
24 h
48 h
72 h
96 h
120 h
time post-seeding
C
48 h
72 h
morphology
(constitutive
DsRed)
Wnt activity
(iRFP)
D
proportion of cells (%)
8
6
4
2
0
t = 48 hrs
t = 72 hrs
$10^0$
$10^1$
$10^2$
$10^3$
$10^4$
$10^5$
iRFP fluorescence (AU)
E
time $t_{fix}$:
84 h
90 h
96 h
102 h
108 h
120 h
144 h
F
morphological
center-of-mass
ΔCOM
signaling activity
center-of-mass

**Supplementary Figure 1.** (**A**) Characterization of off-kinetics of destabilized iRFP reporter of Wnt activity ($P_{TCF/LEF}$-iRFP-PEST). (**B**) Experimental protocol for gastruloid growth with 'LIF only' seed culture media. (**C**) Patterns of Wnt activity in gastruloids grown according to protocol in Figure S1B. Wnt activity is heterogeneous both immediately before (t = 48 h) and immediately after (t = 72) CHIR treatment. (**D**) Definition of iRFP fluorescence threshold above which cells are designated as 'Wnt active' in Figure 1F, based on separation of fluorescence histograms before and after Wnt treatment in 'symmetry-breaking protocol' (Figure 1C). (**E**) Representative images of Wnt activity patterns quantified in Figure 1G. (**F**) Illustration of Wnt polarization metric quantified in Figure 1G.

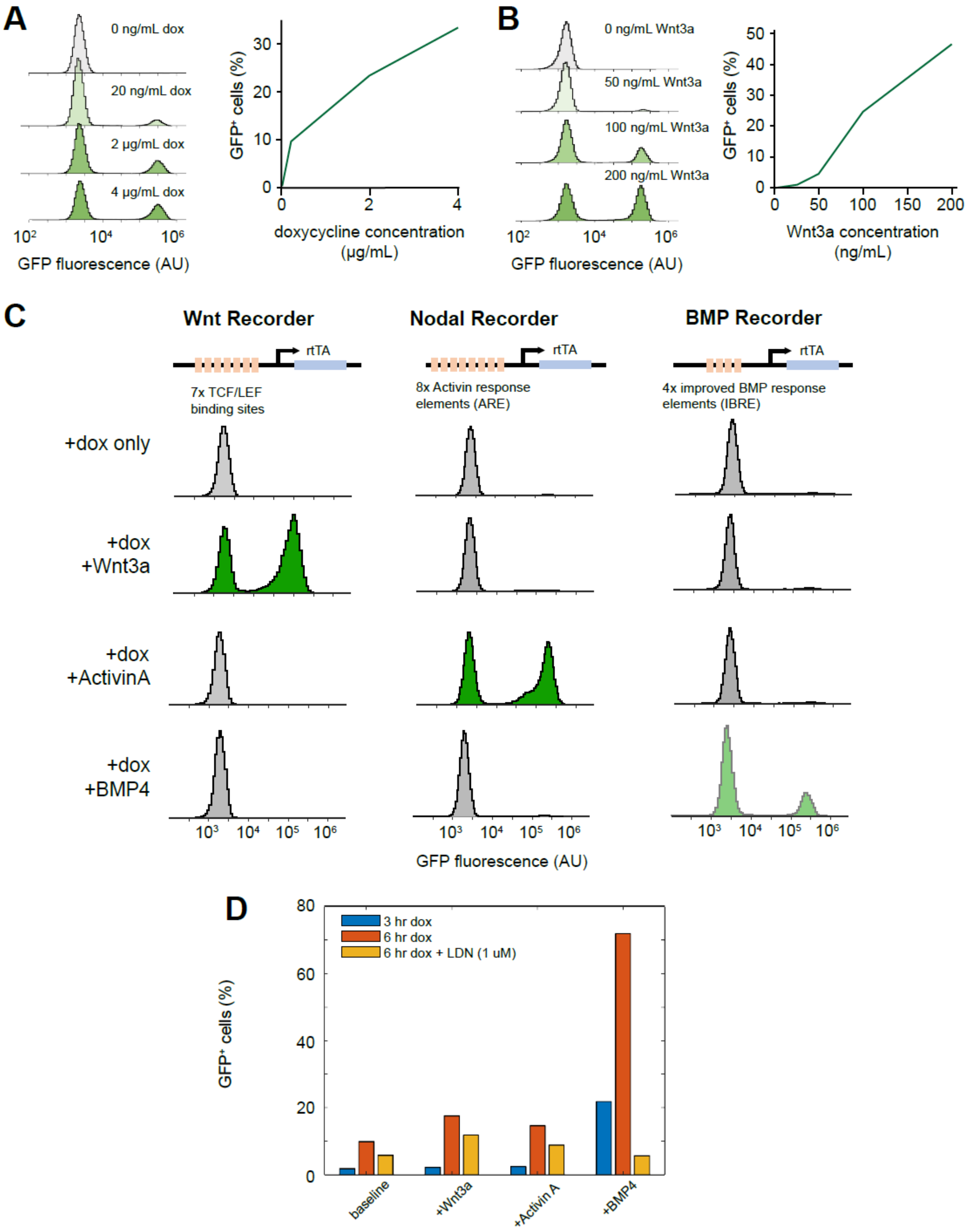


**Supplementary Figure 2: (A)** Recording efficiency in Wnt-Recorder cells depends on doxycycline concentration (3 µM CHIR medium, 1 h recording window). **(B)** Recording efficiency in Wnt-Recorder cells depends on the concentration of recombinant Wnt3a ligand

(200 ng/mL doxycycline, 24 h recording window). **(C)** Flow cytometry histograms corresponding to recorder-ligand crosstalk measurement in Figure 2E. **(D)** BMP-Recorder labeling is sensitive to BMP inhibition with LDN-193189.

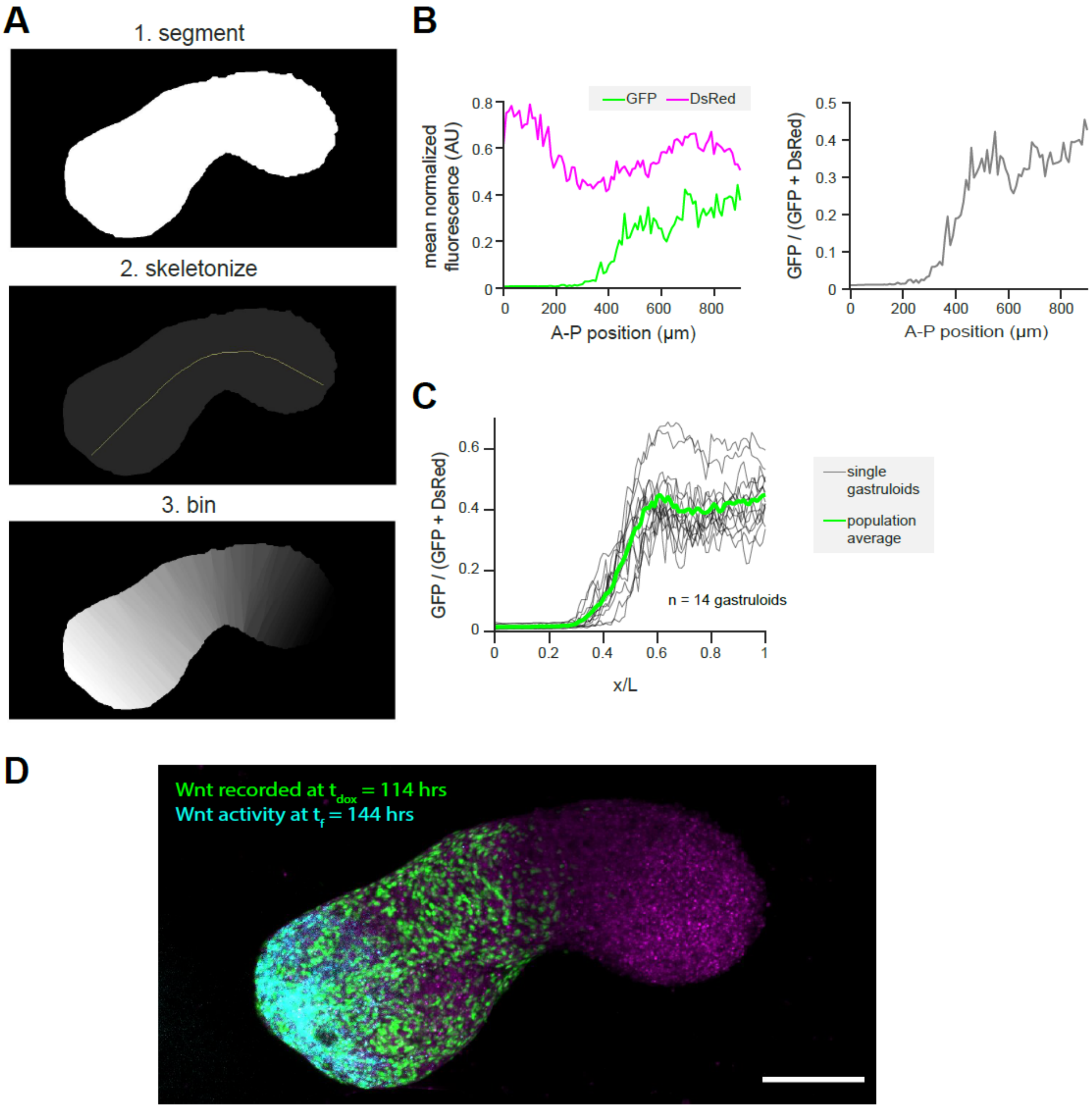


**Supplementary Figure 3: (A)** Illustration of image processing pipleline. Maximum intensity projections of gastruloids are segmented, skeletonized, and then binned to assign an A-P coordinate to each pixel (**Methods**). **(B)** Left: Example quantification of average A-P profiles of both GFP and DsRed expression within a single gastruloid ($t_{dox}$ = 114 h, $t_f$ = 144 h). Pixel intensities were separately normalized relative to maximum image intensities in each channel. Right: relative fraction of GFP labeling across the A-P axis within the same gastruloid (**Methods**). **(C)** Overlay of relative fraction of GFP labeling in individual gastruloids for the same recording condition, along with the mean profile for this condition. **(D)** Simultaneous measurement of the final Wnt activity pattern and recorded Wnt activity signal within the same gastruloid.

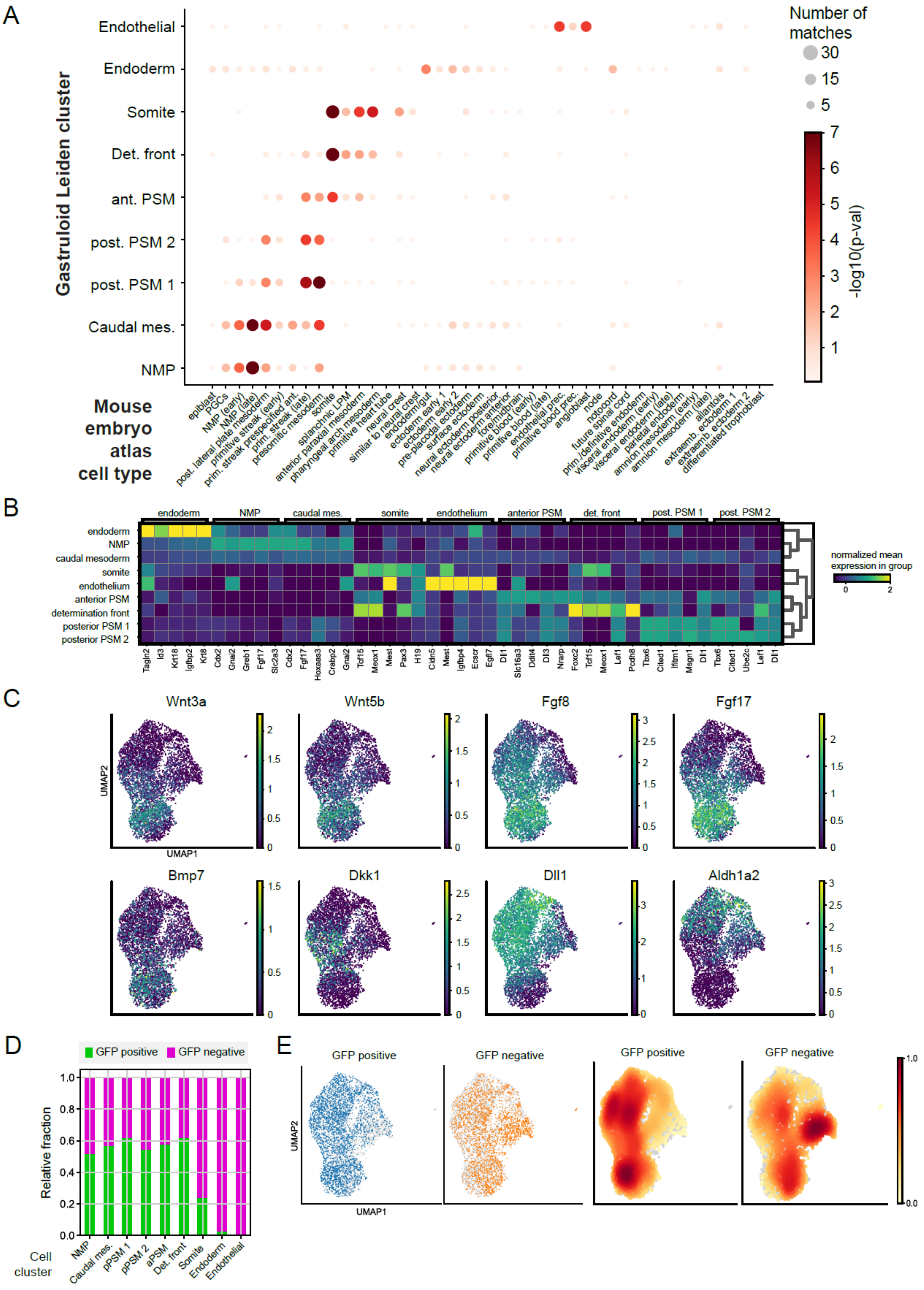
A
Gastruloid Leiden cluster
Endothelial
Endoderm
Somite
Det. front
ant. PSM
post. PSM 2
post. PSM 1
Caudal mes.
NMP
Mouse embryo atlas cell type
epiblast
PGCs
NMP (early)
NMP (late)
post. lateral plate mesoderm
primitive streak (early)
prim. streak prespecified ant.
prim. streak (late)
presomitic mesoderm
somite
splanchnic LPM
anterior paraxial mesoderm
pharyngeal arch mesoderm
primitive heart tube
neural crest
similar to neural crest
endoderm/gut
ectoderm early 1
ectoderm early 2
pre-placodal ectoderm
surface ectoderm
neural ectoderm posterior
neural ectoderm anterior
fore/midbrain
primitive blood (early)
primitive blood (late)
endothelial prec.
primitive blood prec.
angioblast
node
notocord
future spinal cord
prim./definitive endoderm
visceral endoderm (early)
visceral endoderm (late)
parietal endoderm
amnion mesoderm (early)
amnion mesoderm (late)
allantois
extraemb. ectoderm 1
extraemb. ectoderm 2
differentiated trophoblast
Number of matches
30
15
5
-log10(p-val)
B
endoderm
NMP
caudal mes.
somite
endothelium
anterior PSM
det. front
post. PSM 1
post. PSM 2
caudal mesoderm
determination front
posterior PSM 1
posterior PSM 2
Tagln2 Id3 Krt18 Igfbp2 Krt8 Cdx2 Gnai2 Greb1 Fgf17 Slc2a3 Cdx2 Fgf17 Hoxaas3 Crabp2 Gnai2 Tcf15 Meox1 Mest Pax3 H19 Cldn5 Mest Igfbp4 Ecscr Egfl7 Dll1 Slc16a3 Ddit4 Dll3 Nrarp Foxc2 Tcf15 Meox1 Lef1 Pcdh8 Tbx6 Cited1 Ifitm1 Msgn1 Dll1 Tbx6 Cited1 Ube2c Lef1 Dll1
normalized mean expression in group
C
Wnt3a
Wnt5b
Fgf8
Fgf17
Bmp7
Dkk1
Dll1
Aldh1a2
UMAP1
UMAP2
D
GFP positive
GFP negative
Relative fraction
Cell cluster
NMP
Caudal mes.
pPSM 1
pPSM 2
aPSM
Det. front
Somite
Endoderm
Endothelial
E
GFP positive
GFP negative
UMAP1
UMAP2
1.0
0.0

**Supplementary Figure 4: (A)** Comparison of gene expression between Leiden clusters identified in gastruloids at $t_{seq}$ = 120 h and annotated cell types from a reference atlas gene expression during mouse gastrulation[29]. Size of dots indicates the number of shared marker genes; color indicates the binomial likelihood of having at least this many marker genes if compared to a random choice of marker genes (**Methods**). **(B)** Visualization of the top 5 marker genes in each Leiden cluster, ordered according to z-score. **(C)** Single-cell expression levels of signaling-associated genes with the gastruloid data set. **(D)** Relative contributions of GFP-positive and GFP-negative cells to different Leiden clusters, normalized according to total cells recovered in both conditions (Methods). **(E)** Left, Left-center: individual cells in data set separated according to Wnt-recording condition. Right-center, right: embedding density of cells according to Wnt-recording condition.

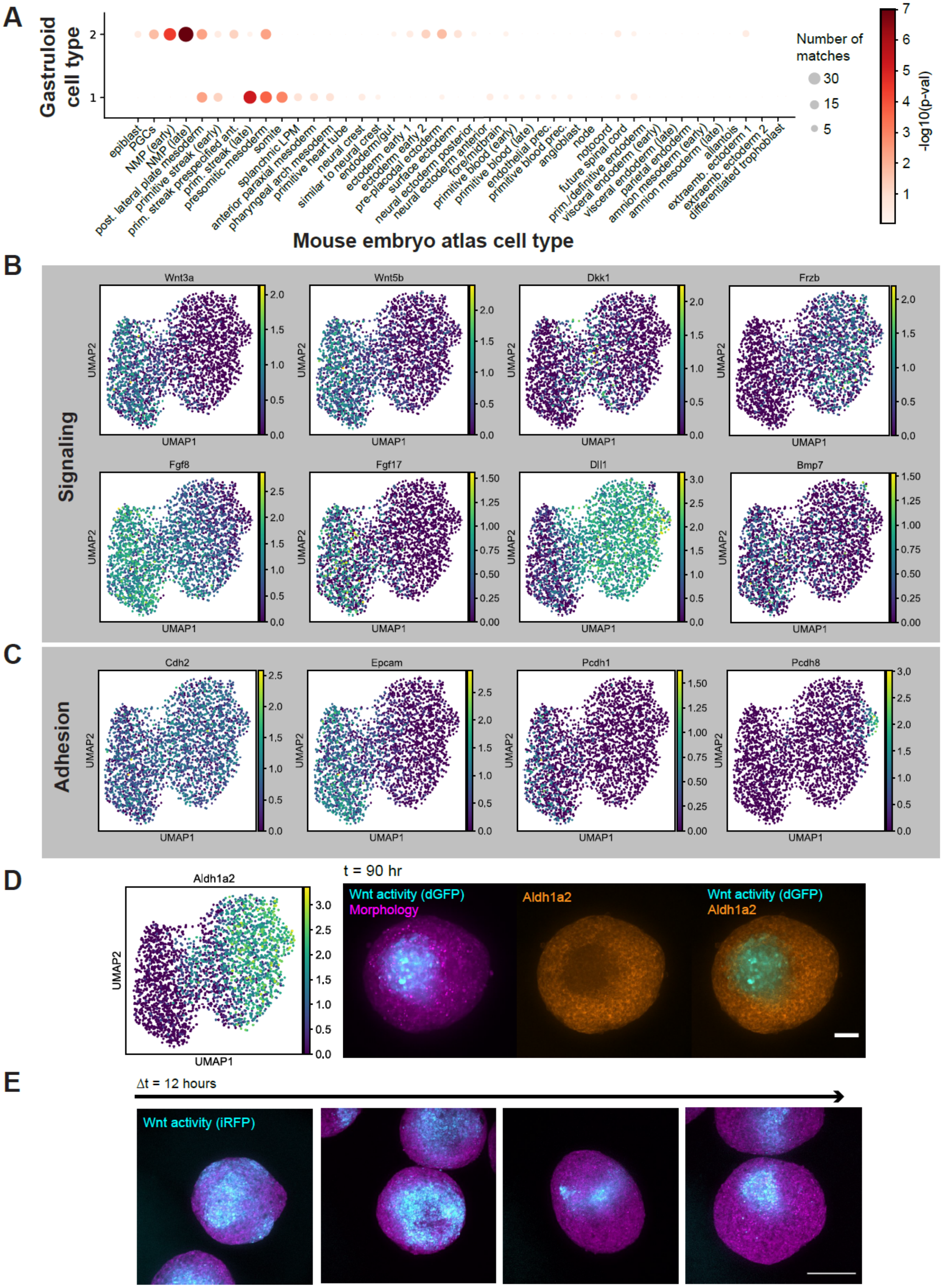
A
Gastruloid cell type
2
1
Number of matches
30
15
5
-log10(p-val)
epiblast
PGCs
NMP (early)
NMP (late)
post. lateral plate mesoderm
primitive streak (early)
prim. streak prespecified ant.
prim. streak (late)
presomitic mesoderm
somite
splanchnic LPM
anterior paraxial mesoderm
pharyngeal arch mesoderm
primitive heart tube
neural crest
similar to neural crest
endoderm/gut
ectoderm early 1
ectoderm early 2
pre-placodal ectoderm
surface ectoderm
neural ectoderm posterior
neural ectoderm anterior
fore/midbrain
primitive blood (early)
primitive blood (late)
endothelial prec.
primitive blood prec.
angioblast
node
notocord
future spinal cord
prim./definitive endoderm
visceral endoderm (early)
visceral endoderm (late)
parietal endoderm
amnion mesoderm (early)
amnion mesoderm (late)
allantois
extraemb. ectoderm 1
extraemb. ectoderm 2
differentiated trophoblast
Mouse embryo atlas cell type
B
Signaling
Wnt3a
Wnt5b
Dkk1
Frzb
Fgf8
Fgf17
Dll1
Bmp7
UMAP1
UMAP2
C
Adhesion
Cdh2
Epcam
Pcdh1
Pcdh8
D
Aldh1a2
t = 90 hr
Wnt activity (dGFP)
Morphology
Aldh1a2
Wnt activity (dGFP)
Aldh1a2
E
Δt = 12 hours
Wnt activity (iRFP)

**Supplementary Figure 5: (A)** Comparison of gene expression between Leiden clusters identified in gastruloids at $t_{seq}$ = 96 h and annotated cell types from a reference atlas gene expression during mouse gastrulation[29] (methods). Size of dots indicates the number of shared marker genes; color indicates the binomial likelihood of having at least this many marker genes if compared to a random choice of marker genes (methods). **(B)** Expression of signaling-associated genes within the scRNAseq dataset at $t_{seq}$ = 96 h. **(C)** Expression of additional cell-cell adhesion associated genes with the scRNAseq dataset at $t_{seq}$ = 96 h. **(D)** Left: Aldh1a2 expression is elevated in Wnt-inactive cells at $t_{seq}$ = 96 h. Right: verification of differential Aldh1a2 expression by immunofluorescence. **(E)** Representative images of fixed samples showing transition of Wnt patterns from heterogeneous domains to a single pole, via ring- and streak-like intermediates. Scale bar = 200 um.

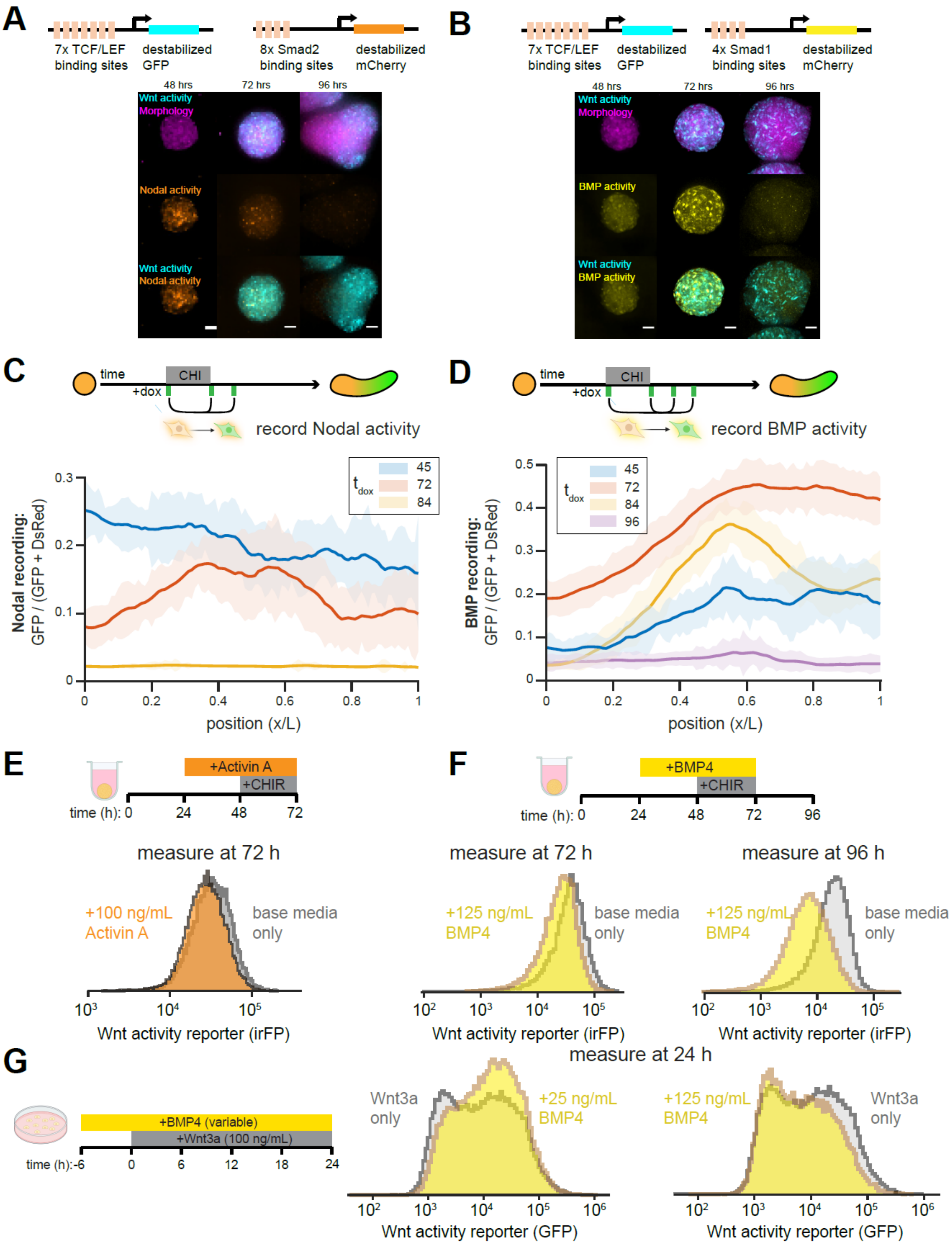
A
7x TCF/LEF binding sites
destabilized GFP
8x Smad2 binding sites
destabilized mCherry
48 hrs
72 hrs
96 hrs
Wnt activity
Morphology
Nodal activity
Wnt activity
Nodal activity
B
7x TCF/LEF binding sites
destabilized GFP
4x Smad1 binding sites
destabilized mCherry
48 hrs
72 hrs
96 hrs
Wnt activity
Morphology
BMP activity
Wnt activity
BMP activity
C
time
CHI
+dox
record Nodal activity
Nodal recording: GFP / (GFP + DsRed)
t dox
45
72
84
position (x/L)
D
time
CHI
+dox
record BMP activity
BMP recording: GFP / (GFP + DsRed)
t dox
45
72
84
96
position (x/L)
E
+Activin A
+CHIR
time (h): 0 24 48 72
measure at 72 h
+100 ng/mL Activin A
base media only
Wnt activity reporter (irFP)
F
+BMP4
+CHIR
time (h): 0 24 48 72 96
measure at 72 h
measure at 96 h
+125 ng/mL BMP4
base media only
+125 ng/mL BMP4
base media only
Wnt activity reporter (irFP)
Wnt activity reporter (irFP)
G
+BMP4 (variable)
+Wnt3a (100 ng/mL)
time (h):-6 0 6 12 18 24
measure at 24 h
Wnt3a only
+25 ng/mL BMP4
+125 ng/mL BMP4
Wnt3a only
Wnt activity reporter (GFP)
Wnt activity reporter (GFP)

**Supplementary Figure 6: (A)** Simultaneous measurement of Wnt activity ($P_{TCF/LEF}$-GFP-PEST) and Nodal activity ($P_{AR8}$-mCherry-PEST) in the same gastruloid with a clonal dual-reporter line. **(B)** Simulatenous measurement of Wnt activity ($P_{TCF/LEF}$-GFP-PEST) and BMP activity ($P_{IBRE4}$-mCherry-PEST) in the same gastruloid with a clonal dual-reporter line. Morphology in (A) and (B) measured with constitutive $P_{CMV}$-TagBFP. Scale bar = 50 µm. **(C)** Recordings of early Nodal activity traced to $t_F$ = 120 h. **(D)** Recordings of early BMP activity traced to $t_F$ = 120 h. **(E)** Flow cytometry measurements of single-cell Wnt activity levels with and without Activin A pretreatment, measured at t = 72 h. **(F)** Flow cytometry measurements of single-cell Wnt activity levels with and without BMP4 pretreatment, measured at t = 72 h (left) and t = 96 h (right). **(G)** Co-treatment of adherent Wnt reporter cells with BMP4 affects responses to Wnt3a treatment. Whereas moderate BMP4 (25 ng/mL) treatment facilitated the Wnt response (middle), higher BMP4 (125 ng/mL) treatment suppressed the Wnt response.

### Supplementary Movie Legends

**Movie S1:** Live imaging of cellular rearrangements during formation and coalescence of Wnt-correlated domains in one gastruloid, separated by fluorescent channel. Scale bar = 100 µm.

**Movie S2:** Live imaging of cellular rearrangements during formation and coalescence of Wnt-correlated domains in multiple gastruloids. Arrows indicate emergence of a Wnt-excluded anterior domain. Scale bar = 100 µm.
